# Ser14-RPN6 Phosphorylation Mediates the Activation of 26S Proteasomes by cAMP and Protects against Cardiac Proteotoxic Stress in Mice

**DOI:** 10.1101/2023.04.05.535705

**Authors:** Liuqing Yang, Nirmal Parajuli, Penglong Wu, Jinbao Liu, Xuejun Wang

## Abstract

**Background:** A better understanding of the regulation of proteasome activities can facilitate the search for new therapeutic strategies. A cell culture study shows that cAMP-dependent protein kinase (PKA) activates the 26S proteasome by phosphorylating Ser14 of RPN6 (pS14-RPN6), but this discovery and its physiological significance remain to be established *in vivo*.

**Methods:** Male and female mice with Ser14 of Rpn6 mutated to Ala (S14A) or Asp (S14D) to respectively block or mimic pS14-Rpn6 were created and used along with cells derived from them. cAMP/PKA were manipulated pharmacologically. Ubiquitin-proteasome system (UPS) functioning was evaluated with the GFPdgn reporter mouse and proteasomal activity assays. Impact of S14A and S14D on proteotoxicity was tested in mice and cardiomyocytes overexpressing the misfolded protein R120G-CryAB (R120G).

**Results:** PKA activation increased pS14-Rpn6 and 26S proteasome activities in wild-type (WT) but not S14A embryonic fibroblasts (MEFs), adult cardiomyocytes (AMCMs), and mouse hearts. Basal 26S proteasome activities were significantly greater in S14D myocardium and AMCMs than in WT counterparts. S14D::GFPdgn mice displayed significantly lower myocardial GFPdgn protein but not mRNA levels than GFPdgn mice. In R120G mice, a classic model of cardiac proteotoxicity, basal myocardial pS14-Rpn6 was significantly lower compared with non-transgenic littermates, which was not always associated with reduction of other phosphorylated PKA substrates. Cultured S14D neonatal cardiomyocytes displayed significantly faster proteasomal degradation of R120G than WT neonatal cardiomyocytes. Compared with R120G mice, S14D/S14D::R120G mice showed significantly greater myocardial proteasome activities, lower levels of total and K48-linked ubiquitin conjugates and of aberrant CryAB protein aggregates, less reactivation of fetal genes and cardiac hypertrophy, and delays in cardiac malfunction.

**Conclusions:** This study establishes in animals that pS14-Rpn6 mediates the activation of 26S proteasomes by PKA and that the reduced pS14-Rpn6 is a key pathogenic factor in cardiac proteinopathy, thereby identifies a new therapeutic target to reduce cardiac proteotoxicity.

## INTRODUCTION

The ubiquitin-proteasome system (**UPS**) is responsible for the degradation of most intracellular proteins. By targeted and timely degradation of terminally misfolded proteins, the UPS is pivotal to protein quality control (**PQC**) which senses and minimizes the level and toxicity of misfolded proteins.^1^ UPS-mediated proteolysis generally consists of two sequential steps: covalent attachment of a substrate protein to a chain of ubiquitin molecules, predominantly through K48 linkages, and subsequent degradation of the ubiquitinated protein by the 26S proteasome.^2^ It was widely assumed that the rate of UPS-mediated protein degradation is solely determined by the rate of ubiquitination. However, emerging evidence suggests that the functionality of 26S proteasomes is vigorously regulated and dedicates the degradation efficiency of ubiquitinated proteins, especially misfolded proteins.^3–5^ Therefore, a better understanding of the regulation of proteasome functioning should facilitate the search for ways to accelerate the breakdown of unwanted and toxic proteins in the cell, a conceivable strategy to prevent or more effectively treat disease with increased proteotoxic stress (**IPTS**).^4^

Phosphorylation has emerged as an important post-translational mechanism for the regulation of 26S proteasomes in health and disease. There is growing evidence that proteasome activities can be altered by protein kinases including the cAMP-dependent protein kinase (PKA),^6–8^ protein kinase G (PKG),^9^ dual receptor tyrosine kinase 2 (DYRK2),^10^ and calcium/calmodulin-dependent protein kinase II (CaMKII).^11, 12^ Among these kinases, PKA is the first and arguably the best studied one to phosphorylate proteasomes. It was first reported that PKA activates the proteasome via phosphorylation of Ser120 of RPT6, an AAA-ATPase subunit of the 19S regulatory particle (**RP**) of 26S proteasomes.^6^ However, this is refuted later by a more comprehensive study, which demonstrates exclusively in cultured cells that Ser14 of RPN6 (a non ATPase subunit of the 19S RP), rather than RPT6, is phosphorylated by PKA and mediates PKA-induced proteasome activation.^8^ Our group also has reported that cAMP elevation increases pS14-Rpn6 in a PKA-dependent manner and improves UPS proteolytic function in cultured cardiomyocytes.^13^ Despite compelling *in vitro* evidence, none of the proteasome phosphosites, including Ser14-RPN6, have been genetically tested in animals for their physiological or pathological significance.

Clinical and experimental studies have revealed the occurrence of proteasome functional insufficiency (**PFI**) and cardiac IPTS during the progression from a large subset of heart diseases, including cardiac proteinopathy, to heart failure,^14^ a leading cause of death and disability in humans. Patients with hypertrophic cardiomyopathy or heart failure displayed impaired cardiac proteasome activities.^15^ Myocardial accumulation of ubiquitinated proteins in dilated or ischemic cardiomyopathies and the increase of pre-amyloid oligomers in cardiomyocytes of failing hearts also are indicative of PFI in humans.^16, 17^ With a mouse model of cardiomyocyte-restricted proteasome functional enhancement, PFI was first established as a major pathogenic factor in cardiac proteinopathy and myocardial ischemia/reperfusion (I/R) injury.^18^ Hence, improvement of proteasome function has the potential to become a new therapeutic strategy for the treatment of heart diseases with IPTS.^4^ However, this is hindered by the lack of effective pharmacological methods. Recent work from our lab has provided compelling evidence that inhibition of phosphodiesterase 1 (PDE1), which stimulates both PKA and PKG, facilitates UPS-mediated protein degradation in a PKA- and PKG-dependent manner in cultured cardiomyocytes and effectively attenuates diastolic malfunction and slowed down the progression of the CryAB_R120G_-based model of cardiac IPTS.^13^ The therapeutic benefits of PDE1 inhibition was associated with increased myocardial pS14-Rpn6,^13^ but it remains unknown if Ser14-RPN6 phosphorylation alone can protect against cardiac IPTS in animals.

To address these critical gaps, we have conducted the present study to determine the role of pS14-RPN6 in the activation of 26S proteasomes by PKA in mice and to determine the physiological significance of this phosphorylation in disease progression of the CryAB_R120G_- based proteinopathy mice. Our results provide unequivocal evidence for the first time in animals that pS14-RPN6 is responsible for the activation of 26S proteasomes by PKA. Moreover, we have discovered that myocardial pS14-Rpn6 is selectively decreased during disease progression of the CryAB^R120G^-based proteinopathy; and we demonstrate that this decrease plays a key pathogenic role in the cardiac proteinopathy, thereby identifies elevating pS14- RPN6 as a potential strategy to treat heart disease with IPTS.

## MATERIALS AND METHODS

(An expanded section of Materials and Methods is provided in Online Supplements.)

### Animals

The Rpn6^S14A^ (S14A) and Rpn6^S14D^ (S14D) knock-in mice were created in the C57BL/6J inbred background via a contract to Shanghai Biomodel Organism Science & Technology Development Co., Ltd. (Shanghai, China), using the CRISPR/Cas9 technology to target the point mutation to the endogenous *Psmd11/Rpn6* gene (<u>MGI:1916327</u>) which is located in chromosome 11. These mice had undergone 6 or more generations of back-cross into the C57BL/6J inbred background before they were cross-bred with GFPdgn transgenic (tg) UPS reporter mice or CryAB^R120G^ tg mice,^19, 20^ respectively to assess their impact on UPS performance and cardiac IPTS. The protocols for animal care and use in this study have been approved by University of South Dakota Institutional Animal Use and Care Committee.

### Statistical methods

GraphPad Prism software (San Diego, CA) was used. All continuous variables are presented as Mean±SEM unless indicated otherwise. All data were examined for normality with the Shaprio Wilk’s test prior to application of parametric statistical tests. Unless otherwise indicated, differences between two groups were evaluated by two-tailed unpaired Student’s t test; differences among ≥3 groups were evaluated by one-way or two-way ANOVA followed by Tukey’s test for pairwise comparisons. Serial echocardiographic data were evaluated by two-way repeated measures ANOVA followed by Tukey’s multiple comparisons. A *p* value <0.05 is considered statistically significant.

## RESULTS

### 1. Creation and baseline characterization of Rpn6^S14A^ and Rpn6^S14D^ mice

To facilitate the investigation in to the (patho)physiological significance of pS14-Rpn6, we created the S14A and S14D mice for blockade and mimicry of pS14-Rpn6, respectively. Their genotypes were confirmed by sequencing the targeted segment of the *Psmd11* gene.

Heterozygous and Homozygous S14A and S14D mice are viable and fertile and do not display discernible gross abnormalities compared with their littermate controls during the first 12 months of age, the longest time observed in full cohorts so far. Monthly M-mode echocardiography did not reveal significant difference in LV end-diastolic volume (LVEDV), ejection fraction (EF), fractional shortening (FS), cardiac output (CO), LV mass index (LV Mass/body weight), or body weight (BW) between wild type (WT) and littermate S14A or S14D mice in either male or female cohorts during the first 7 months (S14A) or 6 months (S14D) of age, the oldest cohorts analyzed so far (*data not shown*).

### 2. S14A blocks the proteasome activation by cAMP/PKA in cells and mice

To determine the role of pS14-Rpn6 in the proteasome activation by cAMP/PKA signaling, we first created mouse embryonic fibroblast (MEF) cell lines from WT and homozygous S14A mice and tested their responses to the augmentation of cAMP/PKA signaling by forskolin (Fsk, an adenylate cyclase activator) or piclamilast (Picl, a phosphodiesterase 4 inhibitor). WT MEFs treated with vehicle control displayed a detectable level of pS14-Rpn6; both forskolin and piclamilast induced significant increases in pS14-Rpn6 (**Figure 1A, 1B**). However, pS14-Rpn6 was completely lost in S14A MEFs regardless of the treatments, although similar levels of increases in the phosphorylated forms of other PKA substrates by either treatment were detected in WT and S14A MEFs (**Figure 1C, 1D**). The 26S proteasome chymotrypsin-like peptidase activity was discernibly lower in S14A MEFs than in WT MEFs under basal condition (p<0.005). Treatment with forskolin or both piclamilast and forskolin led to significant increases in 26S proteasome peptidase activities in WT MEFs, but such effect was completely lost in S14A MEFs (**Figure 1E, 1F**). In addition, the increase in the 26S proteasome peptidase activity by forskolin in WT MEFs was abolished by co-treatment with a PKA inhibitor H89 (**Figure 1F**).

**Figure 1.**
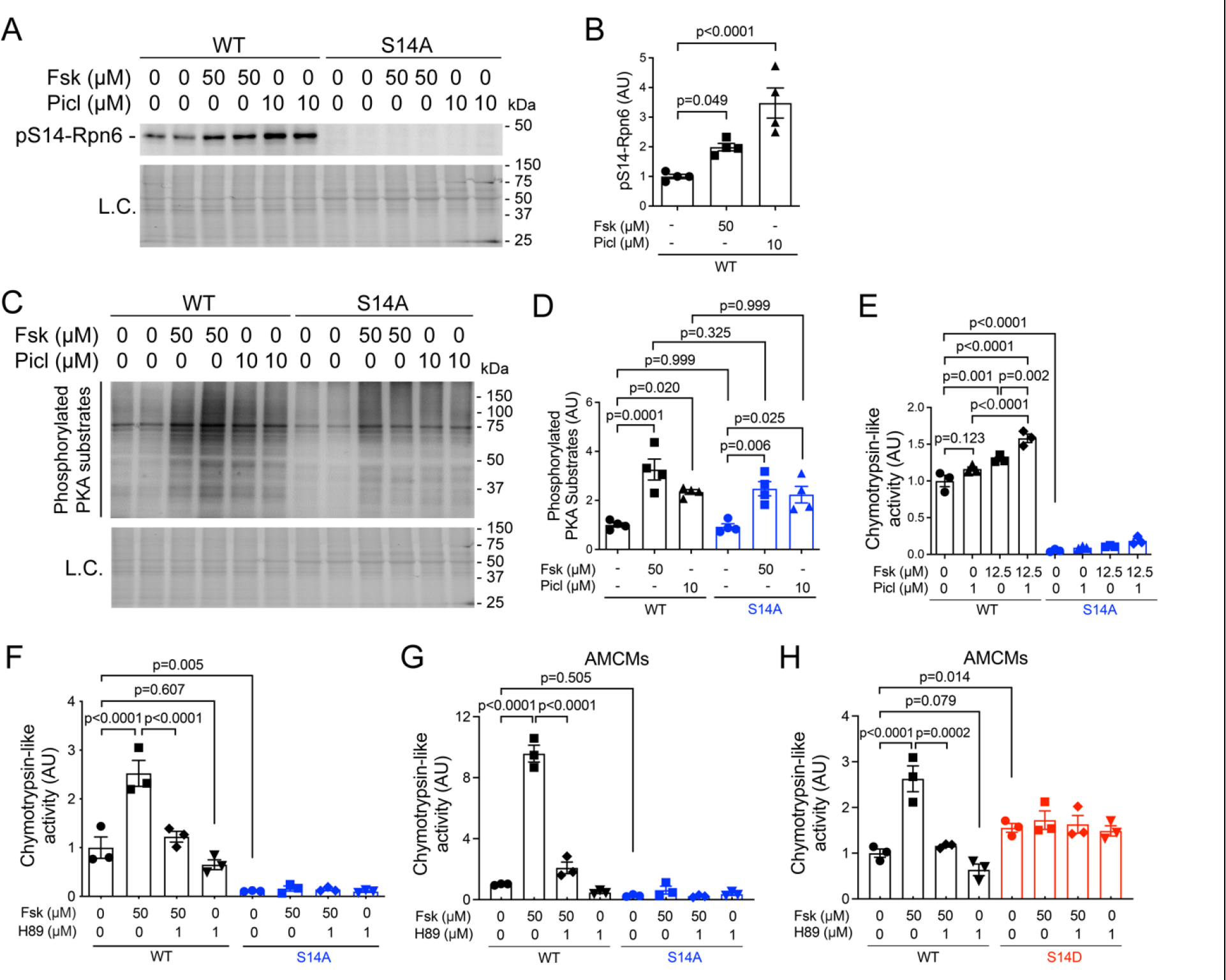
**Ser14-Rpn6 phosphorylation and proteasome activation by cAMP/PKA are lost in cultured cells from S14A mice**. **A** ∼ **F**, Mouse embryonic fibroblasts (MEFs) isolated from wild-type (WT) and homozygous S14A mice were treated with forskolin (Fsk), piclamilast (Picl), or vehicle control (0.1% DMSO) and harvested 3 hours after treatments for crude protein extraction and assays. Representative images (A, C) and pooled densitometry data (B, D) of western blots for Ser14-phosphorylated Rpn6 (pS14-Rpn6) (A and B) and phosphorylated PKA substrates (C and D). L.C., loading control; the in-lane total protein signal from the stain-free image was used to normalize the loading; the same for other figures. Panels E and F show results of 26S proteasome chymotrypsin-like activity assays in MEFs subjected to indicated treatment. **G** and **H**, 26S proteasome chymotrypsin-like activity assays using crude protein extracts of adult mouse cardiomyocytes (AMCMs) from WT, homozygous S14A, or homozygous S14D mice that were cultured and treated with forskolin, H89, both, or vehicle control for 6 hours. . Each dot represents a biological repeat; mean±SEM; two-way ANOVA followed by Tukey’s test.

We also tested the impact of augmentation of cAMP/PKA on proteasome activities in cultured adult mouse cardiomyocytes (AMCMs). The 26S proteasome chymotrypsin-like activity was dramatically elevated in WT AMCMs by forskolin, which was abolished by H89. The basal proteasome peptidase activity was not significantly lower in S14A AMCMs (p=0.505), while it was significantly higher in S14D AMCMs (p=0.014) compared with WT AMCMs. In both S14A and S14D AMCMs, the responses to the treatments were completely lost (**Figure 1G, 1H**).

We then tested the impact of S14A on cAMP/PKA-induced proteasome activation in mice. As expected, pS14-Rpn6 was not detected in S14A mouse myocardium regardless of forskolin treatment (**Figure 2A, 2D**). By contrast, myocardial pS14-Rpn6 in WT mice treated with forskolin (5mg/kg) was 2.5 folds of that in WT mice treated with vehicle control (p<0.0001) and this increase was abolished by pre-treatment of PKA inhibitor H89 (**Figure 2A, 2B**). Myocardial 26S proteasome chymotrypsin-like activities in WT mice treated with forskolin were approximately 3 folds of that in the vehicle control treated WT mice, but this increase was prevented by pre-treatment of H89 (**Figure 2C**). Treatment with forskolin at either 5 mg/kg or 10 mg/kg caused no changes in the proteasome peptidase activities in S14A mice (**Figure 2C**), although they induced comparable levels of phosphorylation of other PKA substrates as in WT mice (**Figure 2D, 2E**), indicative of the requirement of pS14-Rpn6 for PKA to activate 26S proteasomes.

**Figure 2.**
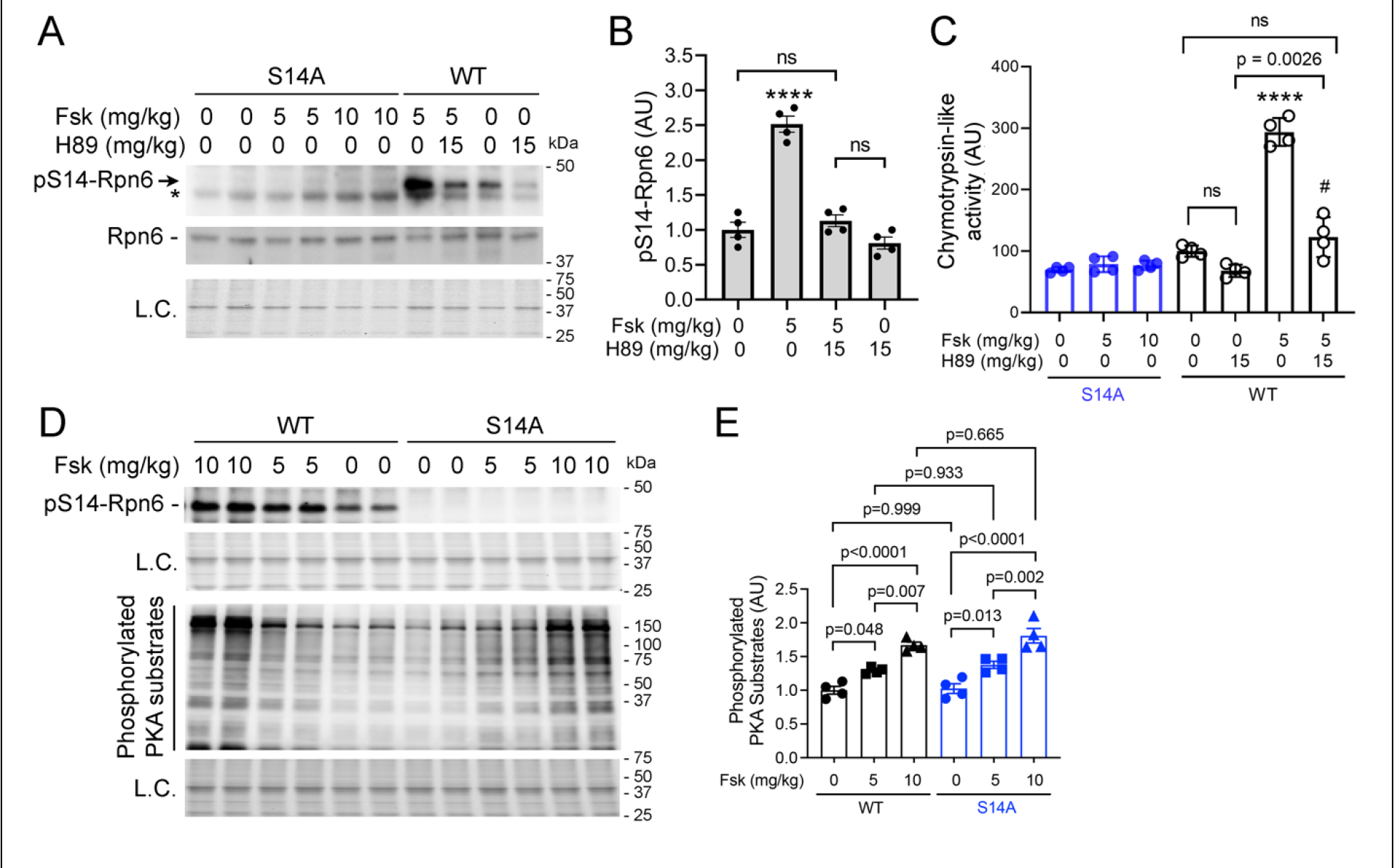
Myocardial Rpn6-Ser14 phosphorylation and proteasome activation by cAMP/PKA signaling are lost in S14A mice. Adult WT and homozygous S14A mice were injected (i.p.) with forskolin, H89, or both and sacrificed 6 hours later to harvest tissues. In the Fsk+H89 group, H89 was injected 15 minutes before forskolin treatment. Total proteins from ventricular myocardium were analyzed. **A** and **B**, Western blot analyses for pS14-Rpn6 (*denotes a nonspecific band) and Rpn6. Shown are representative images from WT and S14A mice (A) and pooled densitometry data from WT mice (B). . **C**, Changes in 26S proteasome chymotrypsin-like peptidase activities. **D** and **E**, Representative images of western blot analyses for the indicated proteins (D) and pooled densitometry data for phosphorylated PKA substrates (E). Each dot represents an individual mouse (male to female 1:1); mean±SEM; two-way ANOVA followed by Tukey’s test. **** p<0.0001 vs. all other groups; # p<0.05 vs. all S14A groups; ns, not significant.

These results together validate in cell cultures that pS14-Rpn6 mediates PKA-induced activation of 26S proteasomes and, more importantly, provide unequivocally the first *in vivo* demonstration that pS14-Rpn6 is required for the activation of 26S proteasomes by cAMP/PKA.

### 3. Myocardial pS14-Rpn6 is selectively decreased in CryAB^R120G^ mice

Since PFI has proven to play a major role in cardiac pathogenesis,^18,21^Next, we determined whether blockage of pS14-Rpn6 would impact on CryAB^R120G^-induced cardiac proteinopathy. CryAB^R120G^ tg (R120G) mice are a well-established model of cardiac proteinopathy, developing concentric cardiac hypertrophy at 3 months of age (3m) and progressing to heart failure by 6m.^20^ We crossbred S14A into R120G mice. To our surprise, pS14-Rpn6 was markedly decreased in R120G mice compared with WT mice at both 3m and 6m (**Figure 3A-3C**). At both ages, Rpn6 protein levels were comparable between R120G and S14A-coupled R120G mice, but both were significantly greater than that in WT mice (**Figure 3A, 3B, 3D**). As a result, the ratios of pS14-Rpn6 to Rpn6 were markedly lower in R120G mice than that in WT mice (**Figure 3E**), which indicate that basal levels of pS14-Rpn6 are significantly decreased in R120G mice during the disease progression. Interestingly, total phosphorylated PKA substrates were decreased at 3m but increased at 6m in R120G mice compared with littermate non-tg controls (**Figure 3F and 3G**), indicating that the decrease in the phosphorylation of Ser14-Rpn6 in R120G mice at 6m is selective. Consistent with decreased pS14-Rpn6 in R120G mice, neither heterozygous nor homozygous S14A exerted discernible effects on cardiac morphometry or function of the R120G mice as revealed by serial echocardiography (**Supplementary Figure S1**). These data together suggest that myocardial pS14-Rpn6 in the R120G mice is so diminished that loss of the residual pS14-Rpn6 does not discernibly alter their proteinopathy progression.

**Figure 3.**
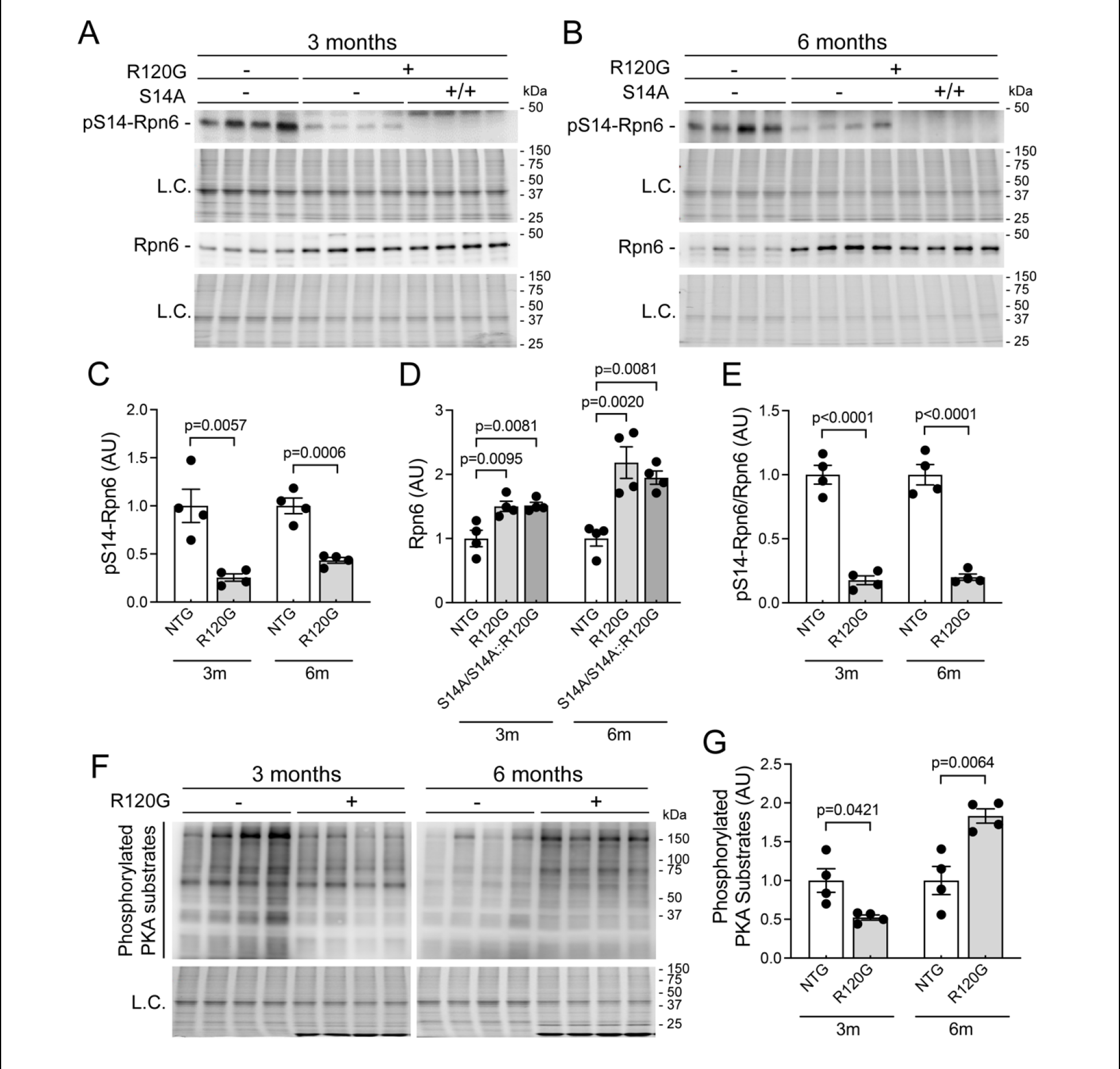
Western blot analyses for myocardial total and Ser14-phosphorylated Rpn6 and phosphorylated PKA substrates. S14A was cross-bred into R120G mice. Adult WT, R120G, and homozygous S14A-coupled R120G mice were sacrificed at 3 months (3m) or 6m of age. Crude protein extracts from ventricular myocardium were used. **A** ∼ **E**, Representative images (A, B) and pooled densitometry data (C ∼ E) of western blot analyses for pS14-Rpn6 and total Rpn6. Homozygous S14A mice were used as the negative control for accurate identification of the pS14-Rpn6 band. **F** and **G**, Representative images (F) and pooled densitometry data (G) of western blot analyses for phosphorylated PKA substrates. Each dot represents an individual mouse (male to female 1:1); mean±SEM; unpaired Student’s *t* test for (C, E, G) and one-way ANOVA followed by Tukey’s test for (D).

### 4. Genetic mimicry of pS14-Rpn6 increases myocardial proteasome activities and enhances UPS performance

As shown in **Figure 1H**, cardiomyocytes from adult S14D mice displayed increased proteasome activities. We next tested the effect of genetic mimicry of pS14-Rpn6 on myocardial proteasomes in mice. GFPdgn protein is a green fluorescent protein (GFP) modified by carboxyl fusion of degron CL1 and has been well-established to inversely reflect UPS performance.^19^ By crossbreeding tg GFPdgn into S14D mice, we found that myocardial GFPdgn protein but not mRNA levels were significantly lower in heterozygous and homozygous S14D-coupled GFPdgn mice, compared with GFPdgn control mice (**Figure 4A-4C**), indicating that S14D decreases myocardial GFPdgn proteins via a post-transcriptional mechanism. The reduction of GFPdgn protein was evident primarily in the cardiomyocyte compartment by confocal microscopy (**Figure 4D**). These data demonstrate that S14D alone can enhance cardiac UPS proteolytic function.

**Figure 4.**
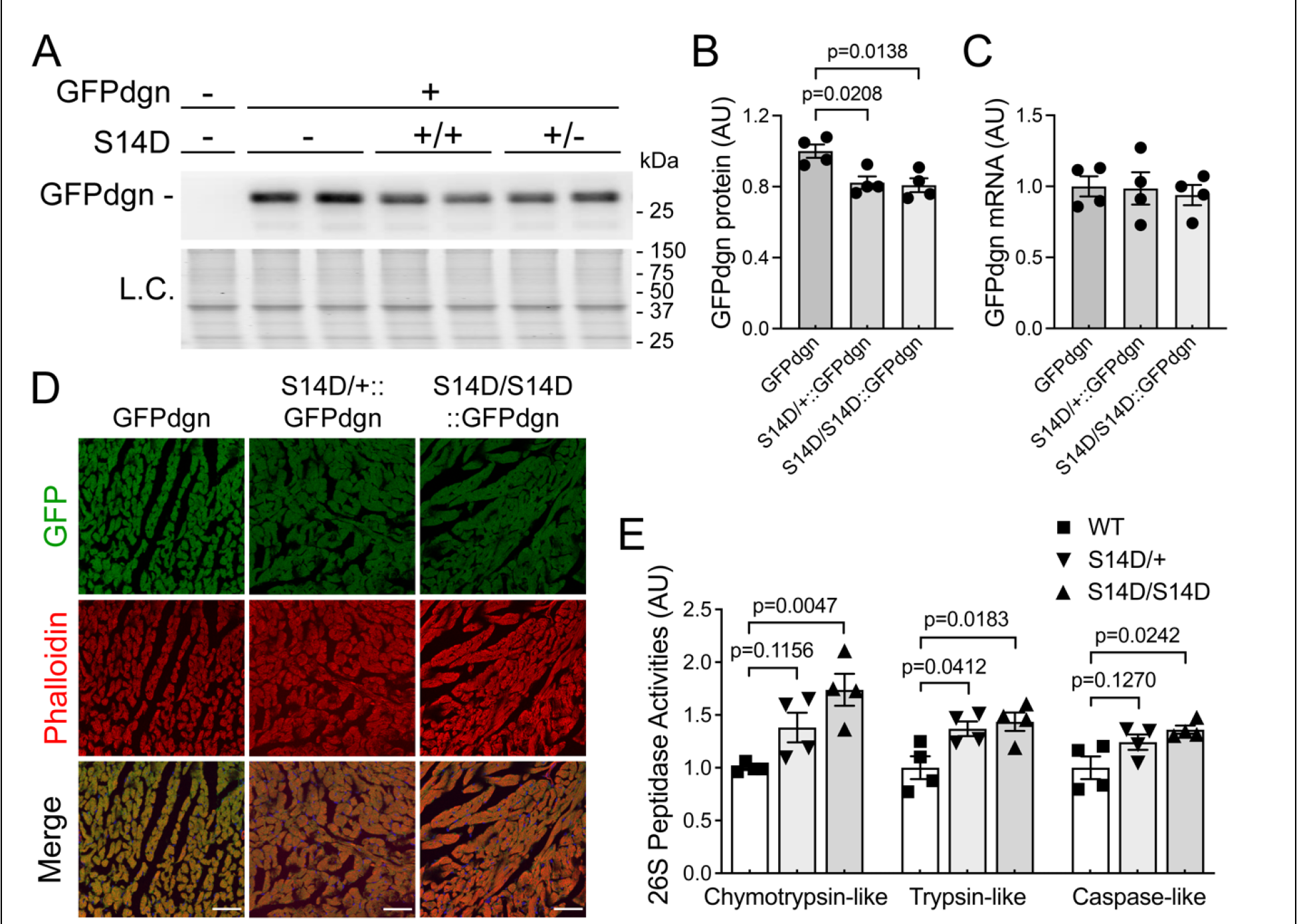
Effect of genetic mimicry of pS14-Rpn6 on myocardial UPS performance and proteasome activities. A. ∼ **D**, Transgenic GFPdgn was cross-bred into S14D mice and the resultant mice were sacrificed at 8 weeks of age. Ventricular myocardium was sampled for protein and RNA extraction or processed for immunofluorescence confocal microscopy. Representative images (A) and pooled densitometry data (B) of western blot for GFPdgn are shown. Myocardial GFPdgn mRNA levels (C) were compared with real-time qPCR analyses. Panel D shows representative confocal micrographs of GFPdgn from direct GFP fluorescence from the indicated groups. Alexa Fluor 568-conjugated Phalloidin was used to stain F-actin to identify cardiomyocytes. Scale bar=50 μm. **E**, Myocardial 26S proteasome peptidase assays. Crude protein extracts from ventricular myocardium of adult mice of the indicated genotypes were used. Each dot represents an individual mouse (male to female 1:1); mean±SEM; one-way ANOVA followed by Tukey’s test

Moreover, we found that myocardial chymotrypsin-like, trypsin-like and caspase-like 26S proteasome activities were increased by ∼75%, ∼50% and ∼40% respectively in homozygous S14D mice (p=0.005, 0.018 and 0.024, respectively), and increased by ∼40%, ∼40% and ∼25% respectively in heterozygous S14D mice (p=0.116, 0.041 and 0.127) compared with that in WT mice (**Figure 4E**), indicating that S14D is capable of increasing all three types of proteasome peptidase activities at baseline.

These results demonstrate that S14D is sufficient to increase myocardial proteasome activities and enhance cardiac UPS performance.

### 5. Genetic mimicry of pS14-Rpn6 increases proteasome activities and facilitates the removal of misfolded proteins in R120G mouse hearts

We next tested whether S14D could enhance proteasome functioning and promote the removal of misfolded CryAB^R120G^ proteins in R120G mouse hearts. By crossbreeding S14D into R120G mice, we first tested myocardial proteasome peptidase activities in the resultant WT, S14D/S14D, R120G, and S14D/S14D::R120G littermates. R120G mice at 1m displayed significantly higher chymotrypsin-like, trypsin-like and caspase-like proteasome activities than WT mice, consistent with a prior report.^21^ Importantly, S14D further increased chymotrypsin-like, trypsin-like and caspase-like proteasome peptidase activities in R120G mice by ∼30%, ∼40% and ∼30%, respectively (**Figure 5A**).

**Figure 5.**
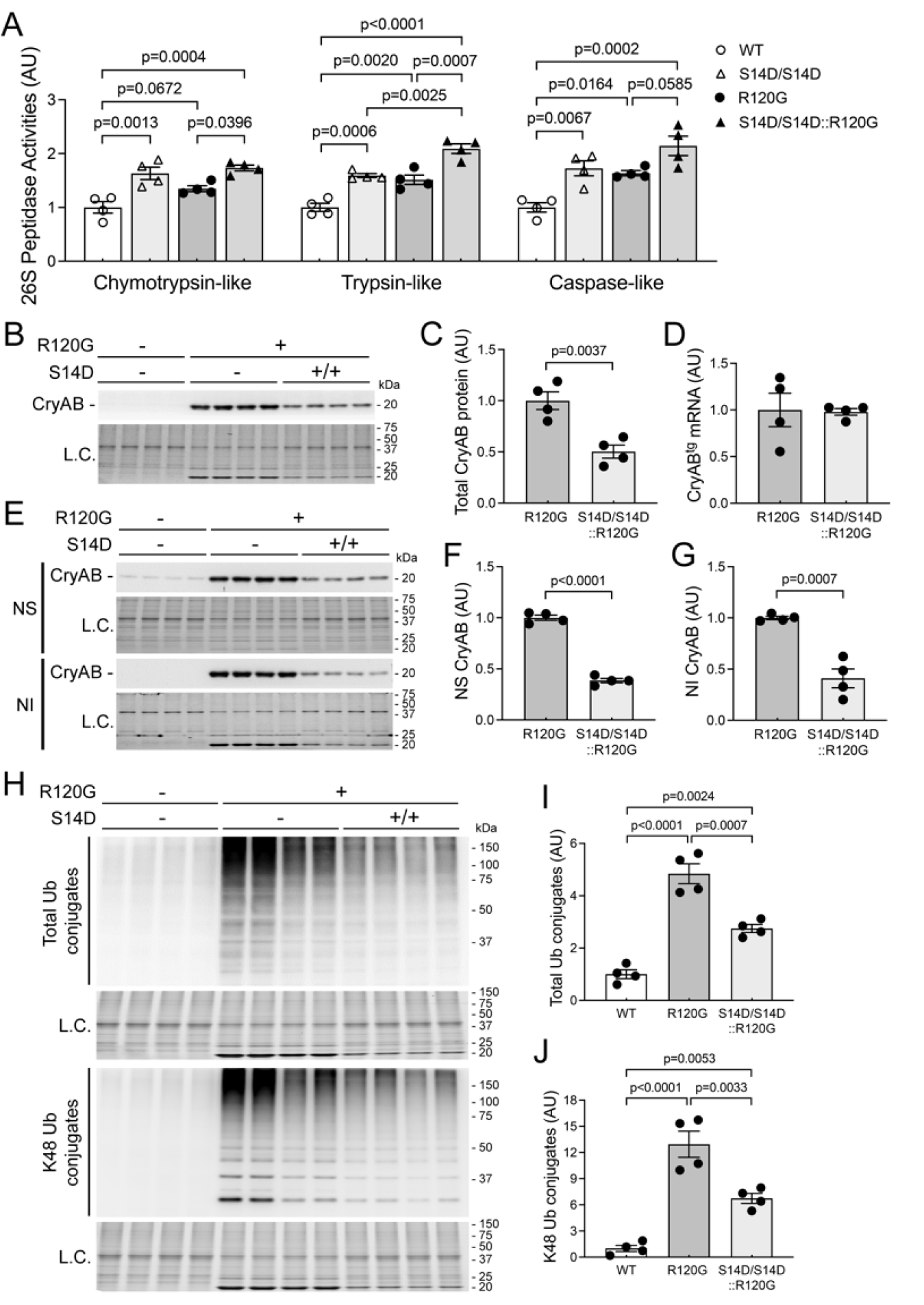
Effect of S14D on myocardial 26S proteasome activities and the abundance of CryAB proteins and ubiquitin conjugates in R120G mice. S14D was cross bred into R120G mice. A, 26S proteasome peptidase activity assays. Crude protein extracts from ventricular myocardium of WT, homozygous S14D, R120G, and S14D/S14D::R120G littermate mice at 1m were used. B ∼ J, Littermate WT, R120G, and S14D/S14D::R120G mice were sacrificed at 6m. Proteins and RNAs from ventricular myocardium were analyzed. Representative images (B, E) and pooled densitometry data (C, F, G) of western blots for CryAB in the total (B, C), NP-40-soluble (NS; F) and insoluble (NI; G) protein fractions are shown. Transgenic CryAB mRNA levels (D) were assessed using real-time reverse-transcriptase PCR. Representative images (H) and pooled densitometry data (I, J) of western blots for total (I) and K48-linked (J) ubiquitin conjugates. Each dot represents an individual mouse (male to female 1:1); mean±SEM; two-way ANOVA followed by Tukey’s test for (A), unpaired Student’s *t* test for (C, D, F, G), and one-way ANOVA followed by Tukey’s test for (I, J).

Myocardial CryAB levels were markedly lower in S14D/S14D::R120G mice compared with R120G control mice at 6m (**Figure 5B, 5C**), without discernible differences in CryAB^R120G^ mRNA levels (p=0.9165, **Figure 5D**). Misfolded proteins undergo aberrant aggregation, forming intermediate oligomers that are highly toxic to the cells. Thus, we examined CryAB abundance in NP-40 soluble (NS) and insoluble (NI; misfolded CryAB oligomers) fractions of myocardial proteins, and found that the protein levels of CryAB in both NS and NI fractions of S14D/S14D::R120G mice were markedly lower than those of the littermate R120G control mice (**Figure 5E-5G**), indicating that degradation of misfolded CryAB by the proteasome in R120G hearts is significantly improved by S14D. Aberrant protein aggregation induced by overexpression of CryAB^R120G^ impairs UPS proteolytic function in the heart, resulting in elevated levels of ubiquitin conjugates.^22^ Both myocardial total and K48-linked ubiquitin conjugates were significantly increased in R120G mice at 6m, and the increase was effectively attenuated when R120G was coupled with S14D (**Figure 5H-5J**).

Taken together, these findings compellingly support the conclusion that genetic mimicry of pS14-Rpn6 increases proteasome proteolytic activity, thereby facilitates the removal of misfolded proteins, and reduces aberrant protein aggregation in the heart, a key pathological process in disease with IPTS.

### 6. Amelioration of CryAB^R120G^-induced cardiac pathology by genetic mimicry of pS14- Rpn6

We next examined aberrant CryAB-positive protein aggregates using immunofluorescence confocal microscopy. At 6m, aberrant CryAB-positive protein aggregates were not detected in WT mouse hearts but were readily detectable in the cardiomyocytes of R120G hearts. More importantly, the CryAB aggregates were clearly less abundant in S14D/S14D::R120G mice compared with their littermate R120G control mice (**Figure 6A**). Consistently, R120G mice developed pronounced cardiac hypertrophy at 6m, as indicated by a higher ventricular weight-to-body weight ratio (VW/BW), compared with that of WT mice (**Figure 6B**). This increase in VW/BW ratios was significantly attenuated in the S14D/S14D::R120G group (**Figure 6B**). At the molecular level, cardiac pathology is commonly accompanied by reactivation of the fetal gene program. Ventricular mRNA levels of atrial natriuretic factor (ANF), brain natriuretic peptide (BNP) and β-myosin heavy chain (Myh7) were markedly upregulated and, reciprocally, α- myosin heavy chain (Myh6) was downregulated in R120G mice; more importantly, these changes were significantly blunted in the S14D/S14D::R120G mice (**Figure 6C-6F**). These results indicate that genetic mimicry of pS14-Rpn6 ameliorates cardiac pathology in proteinopathy animals.

**Figure 6.**
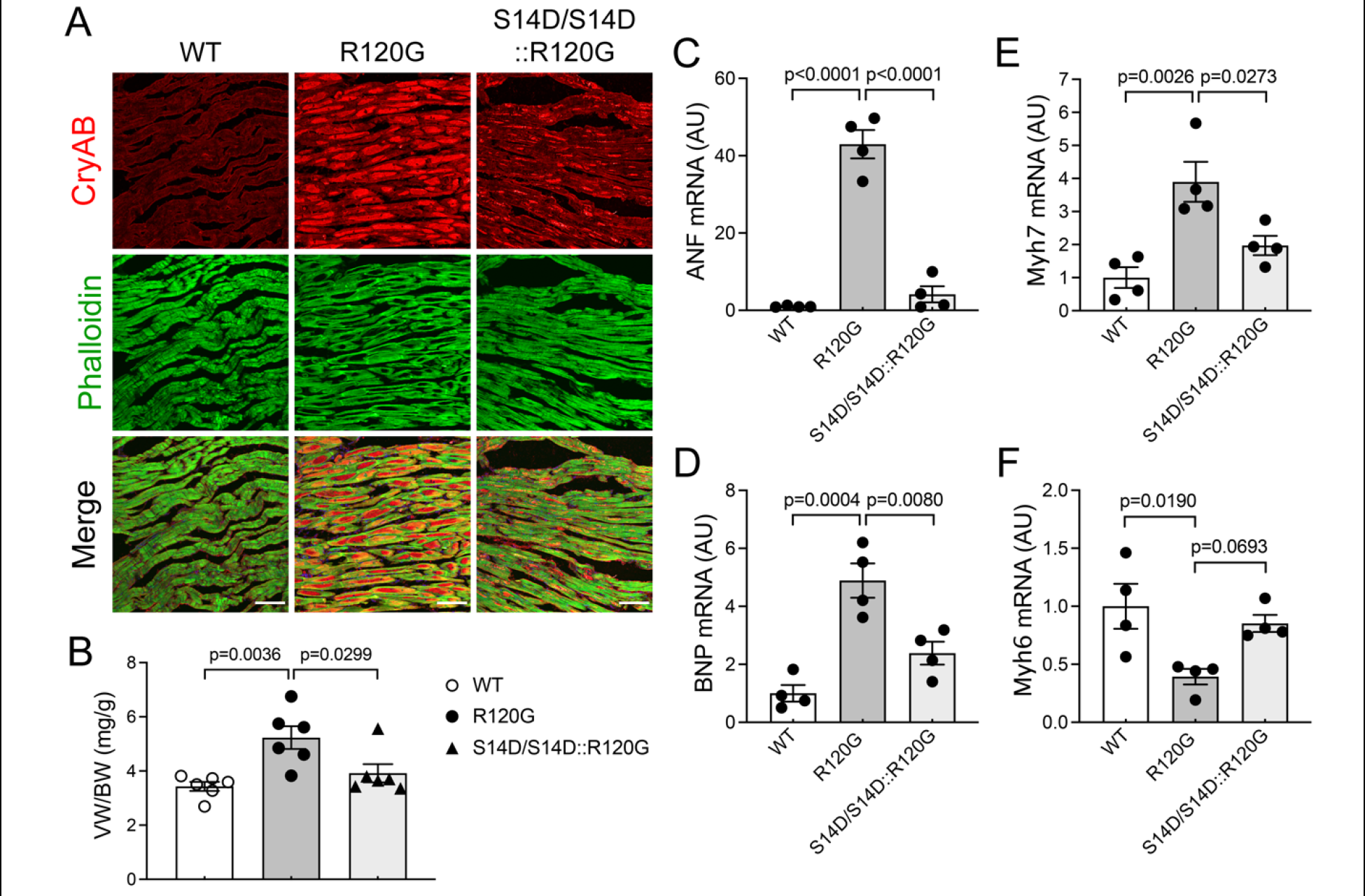
**Genetic mimicry of pS14-Rpn6 ameliorates R120G-induced cardiac pathology. A**, Representative confocal micrographs of WT, R120G, and homozygous S14D-couple R120G mouse ventricular myocardium immunofluorescence stained for CryAB (red) at 6m. Alexa Fluor 488-conjugated Phalloidin was used to stain F-actin to identify cardiomyocytes. Scale bar=100 μm. **B**, Body weight-normalized ventricular weights of mice with indicated genotypes at 6m. **C** ∼ **F**, Comparison of mRNA levels of the indicated genes. Total RNAs extracted from ventricular myocardium were used for real-time qPCR analyses for atrial natriuretic peptide (ANF; C), brain natriuretic peptide (BNP; D), β-myosin heavy chain (Myh7; E), and α-myosin heavy chain (Myh6; F). Each dot represents an individual mouse (male to female 1:1); Mean±SEM; one-way ANOVA followed by Tukey’s test.

### 7. Genetic mimicry of PKA-induced proteasome activation attenuates CryAB^R120G^- induced cardiac malfunction

The results described above establish that S14D suffices to facilitate proteasomal degradation of misfolded proteins and thereby attenuate cardiac proteotoxic stress. This prompted us to further determine the impact on cardiac function. We performed serial echocardiography on WT, homozygous S14D (S14D/S14D), R120G, and S14D/S14D-coupled R120G mice at 1m, 3m, 4.5m, and 6m. All the mice of the same sex had comparable body weights at least during the 6m of observation (**Figure 7A**). Compared with WT mice, R120G mice started displaying a significantly smaller end-diastolic LV internal dimension (LVID;d) and LVEDV, and greater end-diastolic LV posterior wall thickness (LVPW;d), EF and FS, albeit unchanged stroke volume (SV), at 3m (**Supplementary Table S1-S3**), indicative of a compensatory stage. Starting at 3m, R120G mice had discernibly lower heart rate (HR), consistent with prior reports,^13, 20^ and thereby lower CO than WT mice (**Supplementary Table S1-S3**). With the disease progression, the systolic function of R120G mice became compromised, as indicated by the progressive decreases in EF, FS, SV and CO at 4.5m and 6m. These echocardiographic abnormalities were substantially attenuated in the S14D/S14D::R120G groups (**Figure 7B-7E, Supplementary Table S2, S3**). S14D was also trending to, not significantly though, restore the heart rate of R120G mice (**Figure 7F**). These protective effects by S14D knock-in were observed in both female and male mice. These data show that genetic mimicry of pS14-Rpn6 effectively improves cardiac function and attenuates CryAB^R120G^-induced cardiomyopathy.

**Figure 7.**
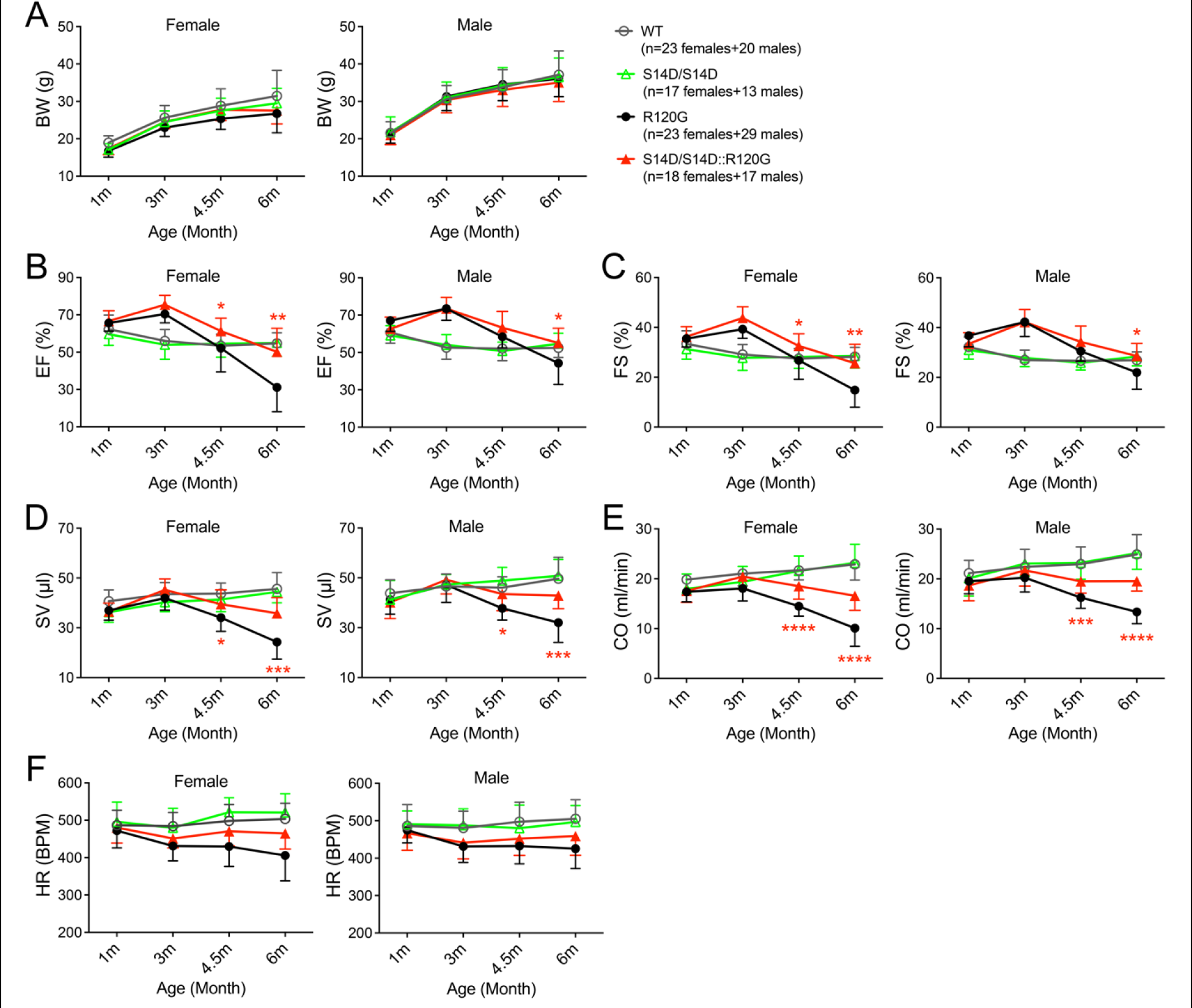
S14D ameliorates R120G-induced cardiac malfunction. Littermate mice of indicated genotypes were subjected to serial echocardiography at 1, 3, 4.5, and 6 months. **A**, Time course of changes in body weight (BW) of females (left panel) and males (right panel). **B** ∼ **F**, Time course of changes in LV functional parameters derived from the serial echocardiography of female (left panels) and male (right panels) littermate mice. Mean±SD; two-way repeated measures ANOVA followed by Tukey’s test; *p<0.05, **p<0.01, ***p<0.001, ****p<0.0001 S14D/S14D::R120G vs. R120G groups; EF, ejection fraction; FS, fractional shortening; SV, stroke volume; CO, cardiac output per minute; HR, heart rate.

Taken together, genetic mimicry of pS14-Rpn6 protects against cardiac proteotoxicity, thereby attenuates CryAB^R120G^-induced cardiac malfunction.

### 8. cAMP augmentation increased proteasome-mediated degradation of misfolded proteins in cardiomyocytes

To test whether the effects observed in the S14D mice are cardiomyocyte-autonomous, we next used cardiomyocyte cultures to examine the effect of cAMP/PKA activation on proteasomal degradation of CryAB^R120G^. Hemagglutinin epitope (HA)-tagged CryAB^R120G^ was overexpressed in cultured neonatal rat cardiomyocytes (NRCMs) via adenoviral gene delivery. NRCMs treated with forskolin exhibited a modest but statistically significant decrease in CryAB^R120G^ protein levels than vehicle treated NRCMs (p=0.0253). This reduction is reversed in the presence of a proteasome inhibitor bortezomib (BZM), suggesting that the lower CryAB^R120G^ protein level is caused by increased proteasome-mediated degradation (**Figure 8A, 8B**). We next measured the proteasome flux by quantifying the difference of CryAB^R120G^ protein levels in the presence or absence of proteasome inhibitor. Forskolin treated NRCMs displayed a dramatically higher proteasome flux of the CryAB^R120G^ proteins than vehicle treated cells (p<0.0001, **Figure 8C**), indicating that cAMP augmentation by forskolin effectively increases proteasome-mediated degradation of misfolded proteins in the cultured cardiomyocytes. This conclusion is also supported by the forskolin-induced decreases in CryAB^R120G^ protein levels in the NP-40 insoluble protein fractions (**Figure 8D, 8E**).

**Figure 8.**
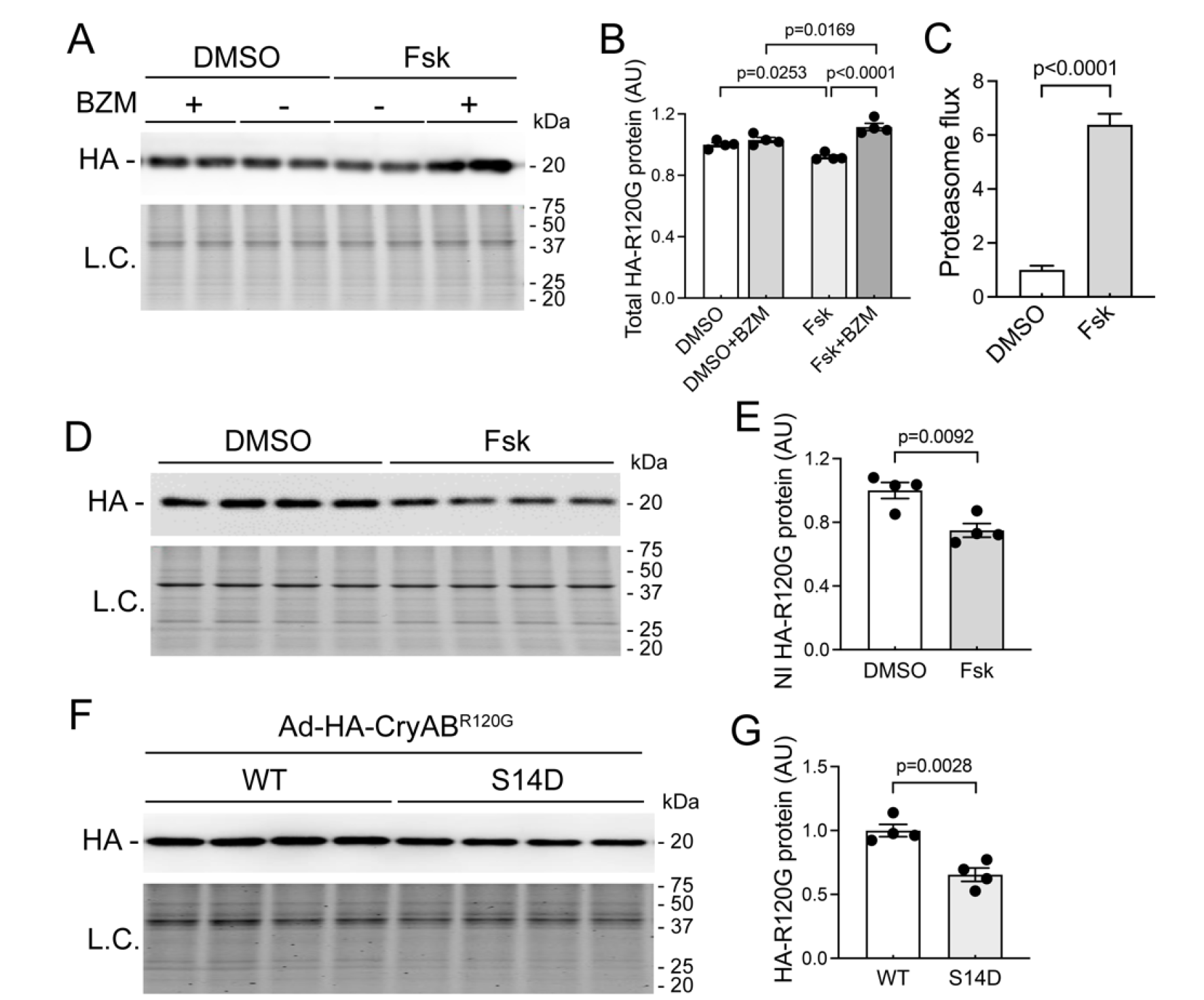
**cAMP augmentation by forskolin (Fsk) or S14D increased proteasome-mediated degradation of CryAB^R120G^ in cardiomyocytes**. **A ∼ C**, Cultured neonatal rat cardiomyocytes (NRCMs) were infected with recombinant adenovirus expressing HA-tagged CryAB^R120G^ (Ad-HA-CryAB^R120G^) in serum-free media for 6 hours followed by 48-hour culture in DMEM containing 2% serum before treatment with forskolin (1 μM) or vehicle control (0.1% DMSO) for 4 hours. The proteasome inhibitor bortezomib (BZM, 50 nM) or volume corrected DMSO were applied to the cultured cells 2 hours after the initiation of Fsk treatment. Representative images (A) and pooled densitometry data (B) of western blot for HA-tagged CryAB, and proteasome flux (C) are presented. The proteasome flux was measured by quantifying the difference of CryAB^R120G^ protein levels in the presence or absence of proteasome inhibition. **D** and **E**, Representative images (D) and pooled densitometry data (E) of western blot for HA in the NP-40-insoluble (NI) protein fraction from NRCMs treated with Fsk (1 μM) or DMSO for 4 hours. **F** and **G,** Representative images (F) and pooled densitometry data (G) of western blot for HA-CryAB^R120G^ in cultured neonatal mouse cardiomyocytes (NMCMs) were infected with Ad-HA-CryAB^R120G^ in serum-free media for 6 hours followed by 48 hour culture in DMEM containing 2% serum before harvest. Whole cell lysates were used for western blots for HA-tagged CryAB. Each dot represents a biological repeat; Mean±SEM; two-way ANOVA followed by Tukey’s test for (B) and unpaired Student’s *t* test for (C, E, G).

Consistent with in vivo data, by overexpressing HA-tagged CryAB^R120G^ proteins in cultured neonatal mouse cardiomyocytes (NMCMs) isolated from WT and homozygous S14D mice, we observed significantly lower CryAB^R120G^ protein levels in S14D/S14D cardiomyocytes than in WT cardiomyocytes (p=0.003, **Figure 8F, 8G**).

These results confirm that both cAMP augmentation and S14D knock-in promote proteasome-mediated degradation of misfolded proteins in cardiomyocytes.

## DISCUSSION

PFI is implicated in the progression from a large subset of heart diseases, including those with IPTS, to heart failure.^9, 14, 18^ There are currently no effective therapies targeting cardiac IPTS.^4^ Hence, a better understanding on proteasome phospho-regulation is undoubtedly of high significance to guide the development of new therapeutic strategies. Previously, Goldberg’s group demonstrated exclusively in cultured cells that cAMP/PKA activates 26S proteasomes by directly phosphorylating RPN6 at Ser14.^8, 23^ Taking advantage of our newly created S14A and S14D mice, the present study has established for the first time in animals that pS14-RPN6 is responsible for the activation of 26S proteasomes by PKA. More importantly, here we have discovered a selective downregulation of myocardial pS14-Rpn6 during cardiac IPTS and further demonstrated that enhancing 26S proteasomes via genetic mimicry of pS14-Rpn6 protects against cardiac IPTS induced by CryAB^R120G^ in animals and thereby effectively curtails cardiac malfunction and disease progression. These new findings establish a novel concept that selective impairment of the proteasome activation by PKA is a key pathogenic factor for cardiac proteinopathy and its targeting can conceivably be exploited to develop new strategies for treating heart disease with IPTS. This is highly significant because a large subset of cardiovascular disease possesses cardiac IPTS.^4^

### 1. pS14-RPN6 is solely responsible for 26S proteasome activation by cAMP/PKA signaling

PKA has been long implicated in proteasome phosphoregulation.^24, 25^ A recent study showed that Ser14-RPN6 was selectively phosphorylated by cAMP/PKA and this phosphorylation was solely responsible for cAMP/PKA-mediated 26S proteasome activation in cultured non-cardiac cells.^8^ We have subsequently demonstrated that cAMP augmentation markedly increased pS14- Rpn6 in cultured cardiomyocytes in a PKA-dependent manner, which was associated with expedited degradation of a surrogate UPS substrate.^13^ However, the physiological relevance of pS14-Rpn6 has not been established in the animals. Here, our experiments in cell cultures and in mice unequivocally establish that Ser14 of Rpn6 is the primary, if not the only, phosphosite responsible for the activation of 26S proteasomes by cAMP/PKA. First, with S14A mice, our findings provide compelling support that pS14-Rpn6 is required for PKA to activate the proteasome. Augmentation of cAMP with an adenylate cyclase activator significantly increased myocardial pS14-Rpn6 proteins and 26S proteasome activities in a PKA-dependent manner in WT mice, but such effects were completely lost in S14A mice (**Figure 2**). Similar findings were observed in cultured MEFs and AMCMs derived from S14A mice (**Figure 1A-1G**). Second, myocardium and cardiomyocytes from S14D mice displayed significantly higher proteasome peptidase activities than WT controls (**Figure 1H, 4E**); and S14D enhanced cardiac UPS proteolytic function (**Figure 4A-4D**), together proving that pS14-Rpn6 is sufficient to activate the proteasome. Along with related prior reports,^8, 23^ the data of the present study suggest that other proteasome phosphosites identified as targets of PKA via *in vitro* assays are unlikely physiologically relevant. Therefore, our newly created knock-in (S14A and S14D) mice confer valuable *in vivo* tools to define the (patho)physiological significance of proteasome activation by cAMP/PKA, adding a new dimension to the (patho)physiology of cAMP/PKA signaling.

### 2. Pathogenic significance of the selective impairment of pS14-RPN6 in cardiac response to IPTS

Desmin-related cardiomyopathies (DRC) are the cardiac aspect of desmin-related myopathies (DRM) that are pathologically featured by intra-sarcoplasmic desmin-positive aberrant protein aggregates.^26, 27^ DRMs arise from mutations in a number of genes, such as desmin,^28^ plectin,^29^ and CryAB.^30^ DRC is the main cause of death in human DRM and exemplifies the pathophysiological significance of IPTS and aberrant protein aggregation in cardiac muscle. The CryAB^R120G^-based DRC mouse model used in the present study recapitulates most aspects of human DRC, including intra-sarcoplasmic aberrant protein aggregation, cardiac hypertrophy, a restrictive cardiomyopathy stage followed by eventually a dilated cardiomyopathy/congestive heart failure stage, and shortened lifespan;^18, 20, 31^ thus the R120G mice are widely used as an animal model to study cardiac IPTS.^13, 32, 33^ PFI is the major pathogenic factor in heart diseases with IPTS, which was best demonstrated by that cardiomyocyte-restricted enhancement of proteasome function markedly decreased aberrant protein aggregation, attenuated cardiac malfunction, and delayed premature death of the R120G mice.^18^ R120G mice display elevated myocardial proteasome peptidase activities to compensate for the misfolded protein overload (**Figure 5A**), consistent with a prior report.^22^ However, to our surprise, pS14-Rpn6 is not part of the natural compensatory response but rather contributes to the insufficiency. This is reflected by marked decrease of myocardial pS14- Rpn6 in R120G mice at both early (3m) and late (6m) stages of disease progression (**Figure 3**). Interestingly, the downregulation of pS14-Rpn6 at 3m was associated with an overall decrease in phosphorylated PKA substrates but, at 6m, it was dissociated with the significant increases of other phosphorylated PKA substrates (**Figure 3F, 3G**), indicating that the decreased pS14- Rpn6 is due to a selective or compartmentalized defect in PKA-mediated phosphorylation of Ser14-RPN6. It will be very interesting and important to delineate the mechanism underlying the decrease of pS14-Rpn6 in cardiac IPTS.

Moreover, we have established the defect in Ser14-RPN6 phosphorylation in the response to IPTS as a major pathogenic factor in cardiac proteotoxicity, and our data provide compelling genetic evidence that enhancement of 26S proteasome functioning by pS14-Rpn6 protects the heart against proteotoxic stress. Upon the adaptive elevations of proteasome activities induced by CryAB^R120G^, S14D further increased myocardial 26S proteasome peptidase activities, which conceivably increases the UPS capacity to degrade misfolded proteins (**Figure 5A**). As a result, the steady-state misfolded CryAB was remarkably reduced (**Figure 5B**). When UPS function becomes inadequate or impaired, misfolded proteins undergo aberrant aggregation with forming intermediate oligomers that is believed highly toxic to the cells.^34^ Here we also show that both total and NP-40-insoluble CryAB (misfolded oligomers) in R120G hearts were markedly decreased by S14D (**Figure 5E**). An increase in total ubiquitinated proteins in the cell can be caused by to increased ubiquitination or inadequate proteasomal degradation and is indicative of defective proteostasis. Consistent with prior reports,^9, 18^ significant increases in myocardial total and K48-linked ubiquitinated proteins, indicative of PFI, were observed in R120G mice, but such increases were substantially attenuated by S14D (**Figure 5H-5J**), demonstrating that genetic mimicry of pS14-Rpn6 effectively facilitates the degradation of ubiquitinated proteins and thereby improves proteostasis. Consistently, aberrant CryAB protein aggregates in R120G mice were markedly decreased as well (**Figure 6A**). At the same time point, the cardiac hypertrophy of R120G mice as indicated by increased VW/BW ratios, was attenuated by S14D, which were further supported by blunted reactivation of representative fetal genes (**Figure 6B- 6F**), indicating that S14D-induced improved removal of pathogenic proteins significantly attenuates cardiac pathology. Functionally, S14D significantly delayed cardiac malfunction, evident by attenuation of the decreases in LV EF, FS, SV, and CO (**Figure 7**). These findings unequivocally demonstrate that genetic mimicry of proteasome activation by PKA protects against cardiac proteotoxicity.

The present study, to our best knowledge, provides the first genetic evidence that PKA-induced proteasome activation protects against proteinopathy in animals, indicating that pS14- RPN6 should be explored as a potentially new strategy to treat heart disease with IPTS.

### 3. Augmentation of pS14-RPN6 as a potentially new strategy to treat heart disease

Despite great advances achieved from numerous preclinical studies over the past decades in the understanding of PKA in the heart, the precise roles of PKA in cardiac pathogenesis and therapeutics remain to be fully understood. A recent study from our group demonstrates that PDE1 inhibition, which activates both PKA and PKG, protects against proteinopathy-based heart failure likely by facilitating proteasome-mediated degradation of misfolded proteins.

R120G mice treated with either a pan PDE1 inhibitor IC86430 or a selective PDE4 inhibitor piclamilast exhibited significantly increased pS14-Rpn6.^13^ Similarly, genetic or pharmacological PDE10A inhibition, which also increases both cAMP and cGMP, was reported to attenuate Ang- II-induced cardiomyocyte hypertrophy *in vitro* and restrain pressure overload-induced cardiac remodeling and dysfunction *in vivo*.^35^ Earlier PDE2 inhibition was also shown to protect hearts from pathological remodeling via a localized cAMP/PKA signaling pathway.^36^ Cardiac hypertrophy is the most common adaptive response of the heart during virtually all heart diseases inherited or acquired, in which increased protein synthesis is inevitable, resulting in increased production of misfolded proteins.^14^ Therefore, it is highly possible that pS14-Rpn6 mediates, at least in part, the attenuation of cardiac hypertrophy by the inhibition of PDE1, PDE2, or PDE10A.

However, controversy exists about the role of PKA in cardiac pathogenesis. Most reports support the involvement of PKA in the development of cardiomyopathy and PKA inhibition as a potential therapeutic target to treat heart disease. For example, activation of cAMP/PKA signaling during the myocardial ischemia contributes to I/R injury;^37^ activation of PKA constitutively^38^ or by chronic sympathetic stimulation^39, 40^ induces maladaptive cardiac hypertrophy and ultimately heart failure; beta blockers are one of the most widely prescribed classes of drug that improve LV function and reverse remodeling in human heart failure.^41^ The present study has demonstrated compellingly that cAMP/PKA protects against cardiac proteotoxicity through improving protein quality control. Therefore, precisely targeting pS14- Rpn6 would potentially serve as an effective therapeutic strategy to treat heart disease with

IPTS, while bypassing possible adverse effects of global PKA activation. Currently, this does not seem to be practical, but a good understanding of the cAMP/PKA signaling at the proteasome nanodomain should facilitate the search for a pharmacological approach to selectively augment cAMP/PKA signaling to the proteasome. An adjunct treatment to minimize undesired effects of PKA activation, such as recently proposed duo-activation of PKA and PKG,^4^ could be an immediately translatable solution.

## Acknowledgments

We thank Megan T. Lewno, Jose R. Lira, and Jack O. Sternburg for their outstanding technical assistance in managing mouse colonies and genotyping for this study.

## Supplemental Material

i. Expanded Materials and Methods^42–46^
ii. Supplemental Tables S1–S3
iii. Supplemental Figures S1 Major Resource Table

## Non-standard Abbreviations and Acronyms

AMCM: adult mouse cardiomyocyte
Fsk: forskolin
IPTS: increased proteotoxic stress
LV: left ventricle/ventricular
MEF: mouse embryonic fibroblast
NRCM/NMCM: neonatal rat/mouse cardiomyocyte
PFI: proteasome functional insufficiency
Picl: piclamilast
PQC: protein quality control
pS14-Rpn6: Serine14-phophorylated Rpn6
S14A: Rpn6/Psmd11^S14A^
S14D: Rpn6/Psmd11^S14D^
UPS: ubiquitin-proteasome system
WT: wild-type

## Novelty and Significance

### What is Known?

- Increased proteotoxic stress (IPTS) and proteasome functional insufficiency (PFI) contribute to cardiac pathogenesis.
- Phosphorylation at Ser14 of proteasome subunit RPN6 (pS14-RPN6) mediates specifically PKA-induced 26S proteasome activation in cultured cells.
- Proteasome enhancement may become a new therapeutic strategy for diseases with IPTS.

### What New Information Does This Article Contribute?

- Establishing in animals that pS14-RPN6 is responsible for the activation of 26S proteasomes by PKA.
- Selective defect in myocardial pS14-RPN6 represents a major pathogenic factor for cardiac proteinopathy.
- Genetic mimicry of pS14-RPN6 increases proteasome activities and proteasomal degradation of misfolded proteins and protects against cardiac proteinopathy in animals.

Means to enhance the proteasome is highly sought. The (patho)physiological significance of PKA was extensively studied but its role in proteostasis remains obscure. A cell culture study reveals that pS14-RPN6 mediates PKA-induced activation of 26S proteasomes but this discovery and its importance remain to be established in vivo. By creating knock-in mice and cells to block or mimic pS14-Rpn6, here we have established for the first time in animals that pS14-RPN6 mediates the activation of 26S proteasomes by cAMP/PKA and that pS14-RPN6 can be augmented to reduce proteotoxicity, thereby identifying potentially a new strategy to treat disease with IPTS.

## References

1. Wang X, Robbins J. Heart failure and protein quality control. Circ Res. 2006;99:1315–1328. doi: 10.1161/01.RES.0000252342.61447.a2

2. Ciechanover A, Schwartz AL. The ubiquitin-proteasome pathway: the complexity and myriad functions of proteins death. Proc Natl Acad Sci U S A. 1998;95:2727–2730. doi: 10.1073/pnas.95.6.2727

3. Collins GA, Goldberg AL. The Logic of the 26S Proteasome. Cell. 2017;169:792–806. doi: 10.1016/j.cell.2017.04.023

4. Wang X, Wang H. Priming the Proteasome to Protect against Proteotoxicity. Trends Mol Med. 2020;26:639–648. doi: 10.1016/j.molmed.2020.02.007

5. Bard JAM, Bashore C, Dong KC, Martin A. The 26S Proteasome Utilizes a Kinetic Gateway to Prioritize Substrate Degradation. Cell. 2019;177:286–298.e215. doi: 10.1016/j.cell.2019.02.031

6. Zhang F, Hu Y, Huang P, Toleman CA, Paterson AJ, Kudlow JE. Proteasome function is regulated by cyclic AMP-dependent protein kinase through phosphorylation of Rpt6. J Biol Chem. 2007;282:22460–22471. doi: 10.1074/jbc.M702439200

7. Asai M, Tsukamoto O, Minamino T, Asanuma H, Fujita M, Asano Y, Takahama H, Sasaki H, Higo S, Asakura M, et al. PKA rapidly enhances proteasome assembly and activity in in vivo canine hearts. J Mol Cell Cardiol. 2009;46:452–462. doi: 10.1016/j.yjmcc.2008.11.001

8. Lokireddy S, Kukushkin NV, Goldberg AL. cAMP-induced phosphorylation of 26S proteasomes on Rpn6/PSMD11 enhances their activity and the degradation of misfolded proteins. Proc Natl Acad Sci U S A. 2015;112:E7176–7185. doi: 10.1073/pnas.1522332112

9. Ranek MJ, Terpstra EJ, Li J, Kass DA, Wang X. Protein kinase g positively regulates proteasome- mediated degradation of misfolded proteins. Circulation. 2013;128:365–376. doi: 10.1161/circulationaha.113.001971

10. Guo X, Wang X, Wang Z, Banerjee S, Yang J, Huang L, Dixon JE. Site-specific proteasome phosphorylation controls cell proliferation and tumorigenesis. Nat Cell Biol. 2016;18:202–212. doi: 10.1038/ncb3289

11. Djakovic SN, Schwarz LA, Barylko B, DeMartino GN, Patrick GN. Regulation of the proteasome by neuronal activity and calcium/calmodulin-dependent protein kinase II. J Biol Chem. 2009;284:26655–26665. doi: 10.1074/jbc.M109.021956

12. Jarome TJ, Kwapis JL, Ruenzel WL, Helmstetter FJ. CaMKII, but not protein kinase A, regulates Rpt6 phosphorylation and proteasome activity during the formation of long-term memories. Front Behav Neurosci. 2013;7:115. doi: 10.3389/fnbeh.2013.00115

13. Zhang H, Pan B, Wu P, Parajuli N, Rekhter MD, Goldberg AL, Wang X. PDE1 inhibition facilitates proteasomal degradation of misfolded proteins and protects against cardiac proteinopathy. Sci Adv. 2019;5:eaaw5870. doi: 10.1126/sciadv.aaw5870

14. Wang X, Robbins J. Proteasomal and lysosomal protein degradation and heart disease. J Mol Cell Cardiol. 2014;71:16–24. doi: 10.1016/j.yjmcc.2013.11.006

15. Predmore JM, Wang P, Davis F, Bartolone S, Westfall MV, Dyke DB, Pagani F, Powell SR, Day SM. Ubiquitin proteasome dysfunction in human hypertrophic and dilated cardiomyopathies. Circulation. 2010;121:997–1004. doi: 10.1161/circulationaha.109.904557

16. Weekes J, Morrison K, Mullen A, Wait R, Barton P, Dunn MJ. Hyperubiquitination of proteins in dilated cardiomyopathy. Proteomics. 2003;3:208–216. doi: 10.1002/pmic.200390029

17. Sanbe A, Osinska H, Saffitz JE, Glabe CG, Kayed R, Maloyan A, Robbins J. Desmin-related cardiomyopathy in transgenic mice: a cardiac amyloidosis. Proc Natl Acad Sci U S A. 2004;101:10132–10136. doi: 10.1073/pnas.0401900101

18. Li J, Horak KM, Su H, Sanbe A, Robbins J, Wang X. Enhancement of proteasomal function protects against cardiac proteinopathy and ischemia/reperfusion injury in mice. J Clin Invest. 2011;121:3689–3700. doi: 10.1172/jci45709

19. Kumarapeli AR, Horak KM, Glasford JW, Li J, Chen Q, Liu J, Zheng H, Wang X. A novel transgenic mouse model reveals deregulation of the ubiquitin-proteasome system in the heart by doxorubicin. Faseb j. 2005;19:2051–2053. doi: 10.1096/fj.05-3973fje

20. Wang X, Osinska H, Klevitsky R, Gerdes AM, Nieman M, Lorenz J, Hewett T, Robbins J. Expression of R120G-alphaB-crystallin causes aberrant desmin and alphaB-crystallin aggregation and cardiomyopathy in mice. Circ Res. 2001;89:84–91. doi: 10.1161/hh1301.092688

21. Wang X, Li J, Zheng H, Su H, Powell SR. Proteasome functional insufficiency in cardiac pathogenesis. Am J Physiol Heart Circ Physiol. 2011;301:H2207–2219. doi: 10.1152/ajpheart.00714.2011

22. Chen Q, Liu JB, Horak KM, Zheng H, Kumarapeli AR, Li J, Li F, Gerdes AM, Wawrousek EF, Wang X. Intrasarcoplasmic amyloidosis impairs proteolytic function of proteasomes in cardiomyocytes by compromising substrate uptake. Circ Res. 2005;97:1018–1026. doi: 10.1161/01.RES.0000189262.92896.0b

23. VerPlank JJS, Lokireddy S, Zhao J, Goldberg AL. 26S Proteasomes are rapidly activated by diverse hormones and physiological states that raise cAMP and cause Rpn6 phosphorylation. Proc Natl Acad Sci U S A. 2019;116:4228–4237. doi: 10.1073/pnas.1809254116

24. Pereira ME, Wilk S. Phosphorylation of the multicatalytic proteinase complex from bovine pituitaries by a copurifying cAMP-dependent protein kinase. Arch Biochem Biophys. 1990;283:68–74. doi: 10.1016/0003-9861(90)90613-4

25. Marambaud P, Wilk S, Checler F. Protein kinase A phosphorylation of the proteasome: a contribution to the alpha-secretase pathway in human cells. J Neurochem. 1996;67:2616–2619. doi: 10.1046/j.1471-4159.1996.67062616.x

26. Goldfarb LG, Dalakas MC. Tragedy in a heartbeat: malfunctioning desmin causes skeletal and cardiac muscle disease. J Clin Invest. 2009;119:1806–1813. doi: 10.1172/jci38027

27. McLendon PM, Robbins J. Desmin-related cardiomyopathy: an unfolding story. Am J Physiol Heart Circ Physiol. 2011;301:H1220–1228. doi: 10.1152/ajpheart.00601.2011

28. Goldfarb LG, Park KY, Cervenáková L, Gorokhova S, Lee HS, Vasconcelos O, Nagle JW, Semino-Mora C, Sivakumar K, Dalakas MC. Missense mutations in desmin associated with familial cardiac and skeletal myopathy. Nat Genet. 1998;19:402–403. doi: 10.1038/1300

29. Winter L, Türk M, Harter PN, Mittelbronn M, Kornblum C, Norwood F, Jungbluth H, Thiel CT, Schlötzer-Schrehardt U, Schröder R. Downstream effects of plectin mutations in epidermolysis bullosa simplex with muscular dystrophy. Acta Neuropathol Commun. 2016;4:44. doi: 10.1186/s40478-016-0314-7

30. Vicart P, Caron A, Guicheney P, Li Z, Prévost MC, Faure A, Chateau D, Chapon F, Tomé F, Dupret JM, et al. A missense mutation in the alphaB-crystallin chaperone gene causes a desmin-related myopathy. Nat Genet. 1998;20:92–95. doi: 10.1038/1765

31. Maloyan A, Osinska H, Lammerding J, Lee RT, Cingolani OH, Kass DA, Lorenz JN, Robbins J. Biochemical and mechanical dysfunction in a mouse model of desmin-related myopathy. Circ Res. 2009;104:1021–1028. doi: 10.1161/circresaha.108.193516

32. Tannous P, Zhu H, Johnstone JL, Shelton JM, Rajasekaran NS, Benjamin IJ, Nguyen L, Gerard RD, Levine B, Rothermel BA, et al. Autophagy is an adaptive response in desmin-related cardiomyopathy. Proc Natl Acad Sci U S A. 2008;105:9745–9750. doi: 10.1073/pnas.0706802105

33. Alam S, Abdullah CS, Aishwarya R, Morshed M, Nitu SS, Miriyala S, Panchatcharam M, Kevil CG, Orr AW, Bhuiyan MS. Dysfunctional Mitochondrial Dynamic and Oxidative Phosphorylation Precedes Cardiac Dysfunction in R120G-αB-Crystallin-Induced Desmin-Related Cardiomyopathy. J Am Heart Assoc. 2020;9:e017195. doi: 10.1161/jaha.120.017195

34. Su H, Wang X. The ubiquitin-proteasome system in cardiac proteinopathy: a quality control perspective. Cardiovasc Res. 2010;85:253–262. doi: 10.1093/cvr/cvp287

35. Chen S, Zhang Y, Lighthouse JK, Mickelsen DM, Wu J, Yao P, Small EM, Yan C. A Novel Role of Cyclic Nucleotide Phosphodiesterase 10A in Pathological Cardiac Remodeling and Dysfunction. Circulation. 2020;141:217–233. doi: 10.1161/circulationaha.119.042178

36. Zoccarato A, Surdo NC, Aronsen JM, Fields LA, Mancuso L, Dodoni G, Stangherlin A, Livie C, Jiang H, Sin YY, et al. Cardiac Hypertrophy Is Inhibited by a Local Pool of cAMP Regulated by Phosphodiesterase 2. Circ Res. 2015;117:707–719. doi: 10.1161/circresaha.114.305892

37. Liu Y, Chen J, Fontes SK, Bautista EN, Cheng Z. Physiological and pathological roles of protein kinase A in the heart. Cardiovasc Res. 2022;118:386–398. doi: 10.1093/cvr/cvab008

38. Antos CL, Frey N, Marx SO, Reiken S, Gaburjakova M, Richardson JA, Marks AR, Olson EN. Dilated cardiomyopathy and sudden death resulting from constitutive activation of protein kinase a. Circ Res. 2001;89:997–1004. doi: 10.1161/hh2301.100003

39. Osadchii OE. Cardiac hypertrophy induced by sustained beta-adrenoreceptor activation: pathophysiological aspects. Heart Fail Rev. 2007;12:66–86. doi: 10.1007/s10741-007-9007-4

40. Enns LC, Bible KL, Emond MJ, Ladiges WC. Mice lacking the Cβ subunit of PKA are resistant to angiotensin II-induced cardiac hypertrophy and dysfunction. BMC Res Notes. 2010;3:307. doi: 10.1186/1756-0500-3-307

41. Reiken S, Wehrens XH, Vest JA, Barbone A, Klotz S, Mancini D, Burkhoff D, Marks AR. Beta-blockers restore calcium release channel function and improve cardiac muscle performance in human heart failure. Circulation. 2003;107:2459–2466. doi: 10.1161/01.Cir.0000068316.53218.49

42. Ye S, Dhillon S, Ke X, Collins AR, Day IN. An efficient procedure for genotyping single nucleotide polymorphisms. Nucleic Acids Res. 2001;29:E88–88. doi: 10.1093/nar/29.17.e88

43. Medrano RF, de Oliveira CA. Guidelines for the tetra-primer ARMS-PCR technique development. Mol Biotechnol. 2014;56:599–608. doi: 10.1007/s12033-014-9734-4

44. Xu J. Preparation, culture, and immortalization of mouse embryonic fibroblasts. Curr Protoc Mol Biol. 2005;Chapter 28:Unit 28.21. doi: 10.1002/0471142727.mb2801s70

45. Ackers-Johnson M, Li PY, Holmes AP, O’Brien SM, Pavlovic D, Foo RS. A Simplified, Langendorff-Free Method for Concomitant Isolation of Viable Cardiac Myocytes and Nonmyocytes From the Adult Mouse Heart. Circ Res. 2016;119:909–920. doi: 10.1161/circresaha.116.309202

46. Pan B, Li J, Parajuli N, Tian Z, Wu P, Lewno MT, Zou J, Wang W, Bedford L, Mayer RJ, et al. The Calcineurin-TFEB-p62 Pathway Mediates the Activation of Cardiac Macroautophagy by Proteasomal Malfunction. Circ Res. 2020;127:502–518. doi: 10.1161/circresaha.119.316007

