## Supplemental methods and data for "Ser14-RPN6 Phosphorylation Mediates the Activation of 26S Proteasomes by cAMP and Protects against Cardiac Proteotoxic Stress in Mice"

**To**

### I. Expanded Materials and Methods

#### Animals

The Rpn6$^{S14A}$ (S14A) and Rpn6$^{S14D}$ (S14D) knock-in mice were newly created in the C57BL/6J inbred background via a contract to Shanghai Biomodel Organism Science & Technology Development Co., Ltd. (Shanghai, China), using the CRISPR/Cas9 technology to target the point mutation to the endogenous *Psmd11/Rpn6* gene (MGI:1916327) which is located in chromosome 11. S14A and S14D mice used in the present study were from breeders that had been subjected to at least 6 generations of back-cross into the C57BL/6J inbred background to eliminate any potential off-target mutations resulting from the CRISPR/Cas9 procedures.

The creation and characterization of a transgenic (tg) mouse model expressing GFPdgn were reported before.[1] GFPdgn is a slightly shorter version of GFP$^{u}$, which is an enhanced green fluorescence protein (GFP) modified by carboxyl fusion of degron CL1. GFP$^{u}$ and GFPdgn are both proven substrates for the UPS.[1] GFPdgn tg mice in the C57BL/6J inbred background were used. S14D mice were cross-bred with GFPdgn tg mice respectively to generate heterozygous and homozygous S14D-coupled hemizygous GFPdgn tg (S14D/+::GFPdgn, S14D/S14D::GFPdgn) mice, with their littermates of hemizygous GFPdgn tg mice serving as the control.

The line 134 tg mouse model expressing the R120G-missense mutant αB-crystallin (CryAB$^{R120G}$) was described previously.[2] CryAB$^{R120G}$ tg mice were maintained in the FVB/N inbred background. S14A and S14D mice were cross-bred with CryAB$^{R120G}$ tg mice respectively to generate heterozygous and homozygous S14A- or S14D-coupled hemizygous CryAB$^{R120G}$ tg (S14A/+:: CryAB$^{R120G}$, S14A/S14A:: CryAB$^{R120G}$ or S14D/+:: CryAB$^{R120G}$, S14D/S14D:: CryAB$^{R120G}$) mice, with their littermates of other genotypes used as controls to ensure the resultant mice are in the same genetic background.

The conventional polymerase chain reaction (PCR) was used for detection of transgenic GFPdgn and CryAB$^{R120G}$ in mice using toe or tail clips and specific primers as previously reported.[1, 2] The genotypes of Rnp6$^{S14A}$ and Rpn6$^{S14D}$ mice were determined with a modified PCR strategy using genomic DNA isolated from toe or tail clips, based on the tetra-primer amplification refractory mutation system (ARMS)-PCR principle described previously,[3, 4] in which two non-allele-specific outer primers and two allele-specific inner primers were employed to detect single nucleotide polymorphisms. In brief, each PCR reaction was carried out in a total volume of 20 µl, containing ~200 ng of template DNA, 2 µl of 10X DreamTaq$^{Tm}$ DNA Polymerase Green Buffer, 0.5 U of DreamTaq$^{TM}$ DNA polymerase (#EP0705, Thermo Scientific), 0.2 mM of dNTPs, outer primers, and inner primers; for S14A: 350 nM of outer primers and 150 nM of inner primers; for S14D: 200 nM of outer primers and 200 nM of inner primers. The primer sequences are listed in **Major Resource Tables.** PCR amplifications for both S14A and S14D were performed with the following profile: 94 °C for 3 min, followed by 30 cycles of 30 sec at 94 °C, 30 sec of annealing, and 45 sec at 72 °C, ending with 10 min at 72 °C. Annealing temperature was 74.9 °C and 74.7 °C for S14A and S14D respectively. The PCR products were electrophoresed in 2% agarose gel and then digitally imaged with the Syngene InGenius LHR2 Gel Imaging System. The S14A and S14D genotypes of all mice used in this study were further confirmed through DNA sequencing the PCR products encompassing the mutation.

The protocols for animal care and use in this study have been approved by University of South Dakota Institutional Animal Use and Care Committee (IACUC). The animals were given *ad lib* access to food and water and housed in specific pathogen free control rooms with optimal temperature (22-24°C) and 12-hour light/12-hour dark cycle.

**Mouse embryonic fibroblast (MEF) isolation**

MEFs from wild type (WT) and homozygous S14A mouse embryos were isolated as reported previously.[5] Briefly, pregnant mice at 14.5 days of gestation were anesthetized by injecting 2.5% of Avertin. The abdominal cavity was then opened and the entire uterus containing all embryos was dissected out. Embryos were removed aseptically and placed in sterile phosphate buffer saline (PBS) to wash out blood. After removal of the head and viscera, all fetuses were teased into fine pieces in 25 ml of 0.25 % trypsin- EDTA (#25300-056, Invitrogen) and the tissue was kept in a tube for digestion at 4°C overnight. On the following day, the tubes with half of the trypsin aspirated off were kept in a 37°C water bath to further digest for 30 min. The digested tissue was next mixed with 25 ml of MEF media (Dulbecco's modified Eagle medium, 10% fetal bovine serum, 0.1 mM β-mercaptoethanol, 50 U penicillin, 50 μg/ml streptomycin) and pipetted vigorously, followed by sedimentation by gravity for 1 min. The collected supernatant was plated with additional MEF media for culture in T75 culture flasks. MEFs were amplified and used for experimentation after the sixth passages.

**Isolation and culture of adult mouse cardiomyocytes**

Cardiomyocytes from adult WT, S14A, and S14D mice were isolated as described previously.[6] In brief, chest cavity of a mouse under anesthesia with 2% inhaled isoflurane was opened, and the descending aorta and the inferior vena cava were cut. The heart was perfused via right ventricle with 7 ml of an EDTA buffer (130 mmol/l NaCl, 5 mmol/l KCl, 0.5 mmol/l $NaH_2PO_4$, 10 mmol/l HEPES, 10 mmol/l Glucose, 10 mmol/l BDM, 10 mmol/l Taurine, and 5 mmol/l EDTA) to flush out blood from the heart. The ascending aorta was then clamped. The clamped heart was removed and submerged in a dish of the EDTA buffer and was perfused via the left ventricle with 10 ml EDTA buffer. After that, the heart was transferred to a dish containing a perfusion buffer (130 mmol/l NaCl, 5 mmol/l KCl, 0.5 mmol/l $NaH_2PO_4$, 10 mmol/l HEPES, 10 mmol/l Glucose, 10 mmol/l BDM, 10 mmol/l Taurine, and 1 mmol/l $MgCl_2$) and injected via the left ventricle with 3 ml of the perfusion buffer to clear EDTA. The heart was then transferred to a dish containing a pre-warmed collagenase buffer (0.5 mg/ml Collagenase 2, 0.5 mg/ml Collagenase 4, and 0.05 mg/ml Protease XIV) and subsequently digested by injections of the collagenase buffer. Once the digestion was satisfied, the ventricular tissue was dissociated mechanically, and the enzymatic activity was inhibited using a Stop buffer (perfusion buffer containing 5% FBS). Then, cardiomyocytes were separated and calcium reintroduction buffers (mixture of perfusion buffer and culture media) were introduced in gradual increments by gravity settling. These separated cardiomyocytes were first plated on laminin (15 µg/ml)-coated 6-well dishes in plating media (M199, 5% FBS, 10 mmol/l BDM, and 1x Penicillin /Streptomycin) for 3 hours and cultured in culture media [M199, 0.1% BSA, 1x ITS (insulin, transferrin, selenium), 10 mmol/l BDM, 1x chemically defined lipid concentrate (CD lipid), and 1x Penicillin /Streptomycin] for another 48 hours prior to further experimentation.

**Neonatal rat/mouse cardiomyocyte culture and adenoviral infection**

Neonatal rat cardiomyocytes (NRCMs) were isolated from the ventricles of 2-day-old Sprague-Dawley rats, and neonatal mouse cardiomyocytes (NMCMs) were isolated from the ventricles of 1-day-old WT and S14D mice with the Cellutron Neomyocytes Isolation System (#nc-6031, Cellutron Life Technology) following the manufacturer's instructions, plated on 35 mm dishes in 10% fetal bovine serum, and cultured as describe previously[7]. Adenoviruses harboring the expression cassette for a Hemagglutinin epitope (HA)-tagged CryAB$^{R120G}$ (Ad-HA-CryAB$^{R120G}$) were described.[8] The plated cells were infected with Ad-HA-CryAB$^{R120G}$ virus in serum-free Dulbecco's minimum essential medium (DMEM) for 6 hours. The cells were then cultured in DMEM containing 2% serum for 48 hours before drug treatment/harvest.

**Total protein extraction and western blot analysis**

Total proteins from ventricular myocardium or cultured cardiomyocytes were homogenized in 1x sampling buffer [50mM Tris-HCl (pH 6.8), 2% SDS and 10% glycerol]. 1x complete protease and phosphatase inhibitor cocktail (#HB9105, Hello Bio Inc.) was added to the sampling buffer to protect proteins from endogenous proteases and phosphatases. The protein homogenates were boiled for 5 minutes. After centrifugation at 10,000x g for 10 min at 4°C, the supernatant was collected. Protein concentrations were determined using Pierce™ bicinchoninic acid (BCA) protein assay reagents (#23222 & #23224, Thermo Fisher Scientific). Equal amounts of proteins were fractionated via 10-14% SDS-polyacrylamide gel electrophoresis (SDS-PAGE) and transferred onto a polyvinylidene difluoride (PVDF) membrane using a Trans-Blot apparatus (Bio-Rad, Hercules, CA). After blocking in 2% BSA or 2% ECL™ advanced blocking agents (#RPN418, Cytiva) in TBST for 1 hour at room temperature, the PVDF membrane was immunodetected with primary antibodies and subsequentially appropriate horseradish peroxidase (HRP)-conjugated secondary antibodies (Jackson ImmunoResearch). The secondary antibodies bound to PVDF membrane were detected with SuperSignal™ West Pico PLUS Chemiluminescent Substrate (#34578, Thermo Fisher Scientific). Blots were digitally imaged with the ChemiDoc™ MP imaging system and analyzed with Image Lab software (Bio-Rad, Hercules, CA). The antibodies used are described in **Major Resource Tables**. The in-lane total protein signal derived from the stain-free image was used as the loading control as previously described.[9]

**Extraction of the NP-40–soluble and NP-40–insoluble fractions**

The protocol for the fractionation of NP-40 soluble and insoluble proteins was developed by Guo et al..[10] Briefly, for cultured cells, cells were washed in cold PBS (pH 7.4) and harvested into cell lysis buffer [50 mM Tris-HCl (pH 8.8), 100 mM NaCl, 5 mM $MgCl_2$, 0.5% NP-40, and 2 mM dithiothreitol (DTT)] containing 1x complete protease and phosphatase cocktail. For tissue, frozen ventricular myocardium was powdered and gently homogenized in the cell lysis buffer. After incubation on ice for 30 min, cell lysates were centrifuged at 17,000x g for 15 min at 4°C. The supernatant was collected as the NP-40–soluble (NS) fraction. The NS fraction added with 3x SDS boiling buffer [6% SDS, 20 mM Tris-HCl (pH 8.0), and 150 mM DTT] was boiled for 5 min. The pellets containing NP-40–insoluble (NI) fraction were washed with PBS and resuspended in cell pellet buffer [20 mM Tris-HCl (pH 8.0), 15 mM $MgCl_2$, 2 mM DTT and 1x complete protease and phosphatase inhibitor cocktail], followed by 30-min incubation on ice. The NI fraction was further solubilized in 3x SDS boiling buffer and boiled for 5 min. Protein concentrations were determined with a reducing agent-compatible BCA protein assay kit (#23252, Thermo Fisher Scientific). The NS and NI fractions were later subjected to SDS-PAGE and western blot analysis.

**Proteasome peptidase activity assays**

Proteasome peptidase activity assays were performed as described previously.[11] Briefly, cultured cardiomyocytes or snap-frozen ventricular myocardial tissue were homogenized in the crude protein extraction buffer [50 mM Tris-HCl (pH 7.5), 250 mM sucrose, 5 mM $MgCl_2$, 0.5 mM

EDTA, 1 mM DTT, and 0.025% digitonin] on ice, followed by centrifugation at 6800x g for 10 min at 4°C. The supernatant was collected and protein concentrations were determined with BCA reagents. Five µg of cellular proteins or ten µg of myocardial proteins in 200 µl proteasome assay buffer [50 mM Tris-HCl (pH 7.5), 40 mM KCl, 5 mM $MgCl_2$, 1 mM DTT, and 0.5 mg/ml BSA] were loaded in a 96-well microplate (#655076, Greiner Bio-One, Germany). The chymotrypsin-like, caspase-like and trypsin-like activities of 26S proteasomes were determined in the presence of 0.2 mM, 0.4 mM and 0.4 mM ATP, respectively. The assays were carried out in the presence or absence of the proteasome inhibitors: 0.28 µM bortezomib (BZM, #B-1408, LC Laboratories) for chymotrypsin-like and caspase-like activities, and 5 µM epoxomicin (#A2606, APExBIO) for trypsin-like activity. The proteasome inhibitor suppressible portion of peptide cleavage is attributed to the proteasome. The following fluorogenic substrates were applied: Suc-LLVY-AMC (18 µM; #10008119, Cayman Chemical), Z-LLE-AMC (45 µM; #10008117, Cayman Chemical), and Bz-VGR-AMC (40 µM; #BML-BW9375, Enzo Life Sciences), for chymotrypsin-like, caspase-like and trypsin-like activities, respectively. The temporal changes in fluorescence intensity were measured immediately after the initiation of the reaction by a Perkin Elmer plate reader (Model VICTOR Nivo™, Waltham, MA) at an excitation wavelength of 380 nm and an emission wavelength of 460 nm. The slope of the log phase of the reaction curve was measured as the indicator of the corresponding proteasome activity.

**Echocardiography**

Serial echocardiography was performed on mice as previously reported.[7] In brief, mice were kept in light anesthesia with inhalation of Isoflurane (4% for induction and 1.5% for maintenance) via a face mask. Transthoracic echocardiography was performed using the VisualSonics Vevo 3100 system and a 40-MHz probe (FUJIFILM VisualSonics, Toronto, ON, Canada). A two-dimensional echocardiogram guided M-mode echocardiography was acquired through the left ventricular (LV) anterior and posterior walls at short axis view. Parameters of LV were derived from primary measurements using Vevo LAB software, based on the following formula:

$$\text{Ejection Fraction (EF)} = \frac{\text{LV end-diastolic volume (LVEDV)} - \text{LV End-systolic Volume (LVESV)}}{\text{LVEDV}}$$

$$\text{Fractional Shortening (FS)} = \frac{\text{LV End-diastolic Dimension (LVID;d)} - \text{LV End-systolic Dimension (LVID;s)}}{\text{LVID;d}}$$

$$\text{Stroke Volume (SV)} = \text{LVEDV} - \text{LVESV}$$

$$\text{Cardiac Output (CO)} = \text{SV} \times \text{Heart Rate (HR)}$$

### RNA isolation and qRT-PCR

Total RNA was isolated from ventricular myocardium using the TRI Reagent® (#TR 118, Molecular Research Center Inc.) as described previously.[9] RNA concentrations were determined with a ThermoFisher NanoDrop 2000 UV Spectrophotometer following the manufacturer's instruction. A High Capacity cDNA Reverse Transcription Kit (#4368814, Applied Biosystems) was used to generate cDNA by following the manufacturer's protocol. PowerUp™ SYBR™ Green Master Mix ((#A25742, Applied Biosystems) was used to conduct quantitative real-time PCR (qPCR) analysis. All qPCR reactions were performed under the following condition: 50 °C for 2 min, 95 °C for 10 min, followed by 40 cycles of amplification at 95 °C for 15 sec and 60 °C for 60 sec. All reactions for reference (GAPDH) and target (GFPdgn, $CryAB^{tg}$, ANF, BNP, Myh7 and Myh6) genes were done in duplicate and the average value was used for subsequent quantification. The $2^{-\Delta\Delta Ct}$ method using GAPDH as the normalization gene was used to calculate the relative expression of target genes. Reverse transcription and qPCR were performed with a Techne 5PRIMEG/02 Thermal Cycler and Applied Biosystems StepOnePlus Real-Time PCR system, respectively. The primer sequences used in this study are listed in **Major Resource Tables**.

### Fluorescence staining and confocal microscopy

Immunofluorescence staining and confocal microscopy were performed as described.[7] In brief, ventricular myocardium was fixed with 4% paraformaldehyde and processed for obtaining 7-μm cryosections. For GFPdgn direct fluorescence, the myocardial sections were stained with Alexa Fluor 568-conjugated phalloidin (1:400; #A12380, Invitrogen) to reveal F-actin that identifies cardiomyocytes. For CryAB-positive aggregates, the myocardial sections were sequentially stained with the rabbit anti-CryAB antibody (1:100; #ADI-SPA-223, Enzo Life Sciences), Alexa Fluor 647-conjugated anti-rabbit secondary antibody (1:500; #111-605-003, Jackson ImmunoResearch), and Alexa Fluor 488-conjugated phalloidin (1:2000; #20478, Cayman Chemical). DAPI (#0100-20, SouthernBiotech) was used for staining nuclei. GFPdgn direct fluorescence (green) or CryAB immunofluorescence and the stained F-actin were visualized and imaged using a Leica TCS SP8 confocal microscope (Leica Microsystems Inc., Buffalo Grove, IL). The fluorescence micrographs of the experimental and control groups were collected using the same imaging setting and processed the same way.

### Statistical methods

The GraphPad Prism software (San Diego, CA) was used. All continuous variables are presented as Mean±SEM unless indicated otherwise. All data were examined for normality with the Shaprio Wilk's test prior to application of parametric statistical tests. Tests used for evaluation of statistical significance are specified in figure legends. In brief, unless otherwise indicated, differences between two groups were evaluated by two-tailed unpaired Student's *t* test; differences among 3 or more groups were evaluated by one-way or, where appropriate, two-way ANOVA followed by Tukey's test for pairwise comparisons. Serial echocardiographic data were evaluated by two-way repeated measures ANOVA followed by Tukey's multiple comparisons. A *p* value or adjusted *p* value <0.05 is considered statistically significant.

### II. Supplementary Tables

**Supplementary Table S1**. Parameters derived from mouse echocardiography from WT, R120G, and S14D/S14D::R120G mice at 3 months

| | Female | | | Male | | |
|---|---|---|---|---|---|---|
| | WT (n=23) | R120G (n=23) | S14D/S14D::R120G (n=18) | WT (n=20) | R120G (n=29) | S14D/S14D::R120G (n=17) |
| BW (g) | 25.6±3.3 | 23.0±2.4 | 24.5±1.8 | 30.6±3.7 | 31.3±3.7 | 30.3±3.4 |
| HR (bpm) | 484±37 | 431±40*** | 451±25* | 481±45 | 432±43** | 442±43 |
| EF (%) | 56.07±6.03 | 70.40±4.72**** | 75.46±4.99**** | 52.60±6.22 | 73.53±6.33**** | 73.53±5.98**** |
| FS (%) | 29.10±4.02 | 39.30±3.73**** | 43.75±4.51**** | 26.98±4.02 | 42.30±5.90**** | 42.23±5.03**** |
| SV (μl) | 43.54±4.61 | 41.94±4.89 | 45.22±4.39 | 46.55±4.99 | 47.16±7.05 | 49.21±5.69 |
| CO (ml/min) | 21.05±2.47 | 18.06±2.52** | 20.43±2.54 | 22.42±3.49 | 20.25±2.88** | 21.70±3.13 |
| LVPW;d (mm) | 0.587±0.058 | 0.793±0.076**** | 0.814±0.023**** | 0.618±0.068 | 0.856±0.084**** | 0.813±0.096**** |
| LVPW;s (mm) | 0.918±0.101 | 1.250±0.104**** | 1.311±0.133**** | 0.948±0.104 | 1.359±0.127**** | 1.354±0.158**** |
| LVID;d (mm) | 4.185±0.219 | 3.737±0.209**** | 3.748±0.171**** | 4.427±0.244 | 3.855±0.292**** | 3.926±0.213**** |
| LVID;s (mm) | 2.972±0.286 | 2.272±0.231**** | 2.112±0.232**** | 3.237±0.315 | 2.234±0.335**** | 2.273±0.278**** |
| LVEDV (μl) | 78.25±9.84 | 59.83±8.13**** | 60.14±6.67**** | 89.30±11.60 | 64.66±11.03**** | 67.25±8.20**** |
| LVESV (μl) | 34.71±8.17 | 17.89±4.70**** | 14.92±4.15**** | 42.75±9.77 | 17.50±5.59**** | 18.04±5.43**** |
| LVAW;d (mm) | 0.611±0.075 | 0.825±0.083**** | 0.824±0.081**** | 0.630±0.065 | 0.859±0.074**** | 0.817±0.111**** |
| LVAW;s (mm) | 0.933±0.092 | 1.244±0.109**** | 1.294±0.122**** | 0.942±0.014 | 1.288±0.017**** | 1.297±0.024**** |

Serial echocardiography was performed on mice of indicated genotypes. Parameters at 3 months are presented. Mean±SD, BW, body weight; HR, heart rate; EF, ejection fraction; FS, fractional shortening; SV, stroke volume; CO, cardiac output per minute; LV, left ventricle; LVPW;d, end-diastolic LV posterior wall thickness; LVPW;s, end-systolic LV posterior wall thickness; LVID;d, end-diastolic LV internal dimension; LVID;s, end-systolic LV internal dimension; LVEDV, LV end-diastolic volume; LVESV, LV end-systolic volume; LVAW;d, end-diastolic LV anterior wall thickness; LVAW;s, end-systolic LV anterior wall thickness; two-way repeated measures ANOVA followed by Tukey's test. *p<0.05, **p<0.01, ***p<0.001, ****p<0.0001 vs. WT mice.

**Supplementary Table S2**. Parameters derived from mouse echocardiography from WT, R120G, and S14D/S14D::R120G mice at 4.5 months

| | Female | | | Male | | |
|---|---|---|---|---|---|---|
| | WT (n=23) | R120G (n=23) | S14D/S14D::R120G (n=18) | WT (n=20) | R120G (n=29) | S14D/S14D::R120G (n=17) |
| BW (g) | 28.8±4.5 | 25.3±2.7 | 27.8±2.9 | 33.8±4.7 | 34.5±4.3 | 33.1±4.4 |
| HR (bpm) | 498±44 | 430±53*** | 470±39# | 497±53 | 433±48** | 452±45[#] |
| EF (%) | 53.46±6.14 | 53.14±12.68 | 61.27±6.98**,[#] | 52.10±6.49 | 58.45±5.94* | 63.33±8.70** |
| FS (%) | 27.46±4.01 | 26.80±7.68 | 32.56±4.86*,[#] | 26.68±4.13 | 30.52±3.97 | 34.27±6.38** |
| SV (µl) | 43.72±4.25 | 34.09±5.49**** | 39.48±5.75[#] | 46.09±4.39 | 37.78±4.78**** | 43.53±6.69[#] |
| CO (ml/min) | 21.69±1.93 | 14.48±1.97**** | 18.52±2.63**,[####] | 22.96±3.49 | 16.25±2.16**** | 19.51±2.36*,[###] |
| LVPW;d (mm) | 0.617±0.053 | 0.934±0.097**** | 0.903±0.096**** | 0.648±0.050 | 0.945±0.122**** | 0.898±0.139**** |
| LVPW;s (mm) | 0.939±0.092 | 1.218±0.161**** | 1.283±0.097**** | 0.976±0.093 | 1.276±0.136**** | 1.328±0.181**** |
| LVID;d (mm) | 4.278±0.183 | 3.932±0.328*** | 3.867±0.264**** | 4.430±0.272 | 3.872±0.259**** | 3.978±0.325*** |
| LVID;s (mm) | 3.107±0.268 | 2.899±0.519 | 2.614±0.315**** | 3.256±0.352 | 2.695±0.295*** | 2.627±0.413*** |
| LVEDV (µl) | 82.30±8.25 | 67.92±13.09**** | 65.02±10.16**** | 89.60±12.74 | 65.22±10.25**** | 69.78±13.93** |
| LVESV (µl) | 38.58±7.98 | 33.83±14.46 | 25.54±7.26**** | 43.51±10.92 | 27.44±7.50**** | 26.25±10.09*** |
| LVAW;d (mm) | 0.638±0.054 | 0.932±0.100**** | 0.899±0.061**** | 0.647±0.047 | 0.970±0.115**** | 0.925±0.117**** |
| LVAW;s (mm) | 0.942±0.077 | 1.209±0.164**** | 1.267±0.120**** | 0.949±0.071 | 1.311±0.128**** | 1.313±0.152**** |

Serial echocardiography was performed on mice of indicated genotypes. Parameters at 4.5 months are presented. Mean ± SD; two-way repeated measures ANOVA followed by Tukey's test. *p<0.05, **p<0.01, ***p<0.001, ****p<0.0001 vs. WT mice; $^{\#}$p<0.05, $^{\#\#}$p<0.01, $^{\#\#\#}$p<0.001, $^{\#\#\#\#}$p<0.0001 vs. R120G mice.

**Supplementary Table S3**. Parameters derived from mouse echocardiography from WT, R120G, and S14D/S14D::R120G mice at 6 months

| | Female | | | Male | | |
|---|---|---|---|---|---|---|
| | WT (n=23) | R120G (n=16) | S14D/S14D::R120G (n=15) | WT (n=20) | R120G (n=23) | S14D/S14D::R120G (n=15) |
| BW (g) | 31.5±6.8 | 26.7±5.2 | 27.6±3.6 | 37.1±6.4 | 36.0±4.8 | 35.1±5.1 |
| HR (bpm) | 503±42 | 406±68*** | 465±42 | 505±51 | 425±53*** | 459±51 |
| EF (%) | 54.67±5.70 | 31.15±12.98**** | 50.20±12.61## | 52.49±5.16 | 44.27±11.42* | 55.12±7.88# |
| FS (%) | 28.24±3.73 | 14.84±6.85**** | 25.63±7.61## | 26.93±3.36 | 21.98±6.77* | 28.57±5.07# |
| SV (μl) | 45.64±6.56 | 24.26±6.91**** | 35.81±6.29***,### | 49.67±8.59 | 32.03±7.92**** | 42.88±5.28### |
| CO (ml/min) | 22.89±3.15 | 10.08±3.61**** | 15.58±2.92****,#### | 24.91±3.98 | 13.35±2.36**** | 19.53±1.97***,#### |
| LVPW;d (mm) | 0.650±0.075 | 0.985±0.204**** | 0.989±0.080**** | 0.651±0.059 | 1.047±0.108**** | 0.998±0.116**** |
| LVPW;s (mm) | 0.999±0.097 | 1.145±0.211 | 1.283±0.097**** | 0.980±0.105 | 1.250±0.145**** | 1.327±0.132**** |
| LVID;d (mm) | 4.308±0.239 | 4.084±0.356 | 4.072±0.307 | 4.533±0.285 | 4.293±0.488* | 4.197±0.280* |
| LVID;s (mm) | 3.094±0.263 | 3.492±0.659* | 3.045±0.503 | 3.311±0.249 | 3.298±0.477 | 3.007±0.379 |
| LVEDV (μl) | 83.82±10.76 | 74.35±15.85 | 73.64±12.81 | 94.57±13.62 | 84.24±23.33 | 78.95±12.20* |
| LVESV (μl) | 38.18±7.49 | 53.97±26.00* | 37.83±14.54 | 44.90±8.08 | 46.31±14.78 | 36.07±10.97 |
| LVAW;d (mm) | 0.667±0.056 | 0.950±0.139**** | 0.985±0.076**** | 0.653±0.066 | 1.044±0.126**** | 1.006±0.099**** |
| LVAW;s (mm) | 0.980±0.080 | 1.172±0.172 | 1.274±0.0116****,# | 0.966±0.090 | 1.283±0.135**** | 1.333±0.138**** |

Serial echocardiography was performed on mice of indicated genotypes. Parameters at 6 months are presented. Mean±SD; two-way repeated measures ANOVA followed by Tukey's test. *p<0.05, **p<0.01, ***p<0.001, ****p<0.0001 vs. WT mice; $^{\#}$p<0.05, $^{\#\#}$p<0.01, $^{\#\#\#}$p<0.001, $^{\#\#\#\#}$p<0.0001 vs. R120G mice.

### III. Supplementary Figures

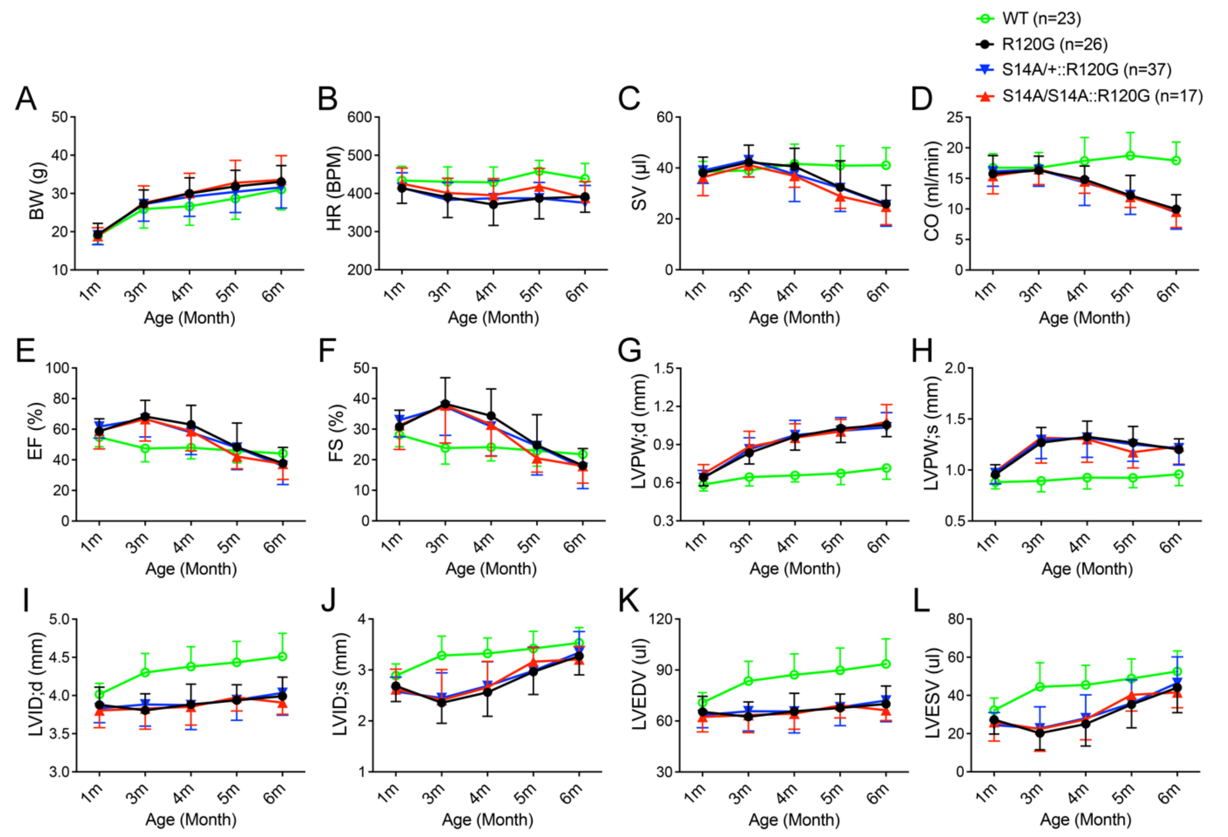


**Figure S1**. **Serial echocardiography for S14AxR120G mice.** Littermate mice of indicated genotypes were subjected to serial echocardiography at 1, 3, 4, 5, and 6 months. LV function parameters derived from the serial echocardiography. The stacked line chart of each panel summarizes the time course of changes in the indicated parameters. Mean±SD; two-way repeated measures ANOVA followed by Tukey's test; no significant differences in any parameters among R120G, S14A/+::R120G, and S14A/S14A::R120G groups. BW, body weight; HR, heart rate; SV, stroke volume; CO, cardiac output per minute; EF, ejection fraction; FS, fractional shortening; LV, left ventricle; LVPW;d, end-diastolic LV posterior wall thickness; LVPW;s, end-systolic LV posterior wall thickness; LVID;d, end-diastolic LV internal dimension; LVID;s, end-systolic LV internal dimension; LVEDV, LV end-diastolic volume; LVESV, LV end-systolic volume.

**References**


1. Kumarapeli AR, Horak KM, Glasford JW, Li J, Chen Q, Liu J, Zheng H and Wang X. A novel transgenic mouse model reveals deregulation of the ubiquitin-proteasome system in the heart by doxorubicin. *Faseb j*. 2005;19:2051-3.

2. Wang X, Osinska H, Klevitsky R, Gerdes AM, Nieman M, Lorenz J, Hewett T and Robbins J. Expression of R120G-alphaB-crystallin causes aberrant desmin and alphaB-crystallin aggregation and cardiomyopathy in mice. *Circ Res*. 2001;89:84-91.

3. Ye S, Dhillon S, Ke X, Collins AR and Day IN. An efficient procedure for genotyping single nucleotide polymorphisms. *Nucleic Acids Res*. 2001;29:E88-8.

4. Medrano RF and de Oliveira CA. Guidelines for the tetra-primer ARMS-PCR technique development. *Mol Biotechnol*. 2014;56:599-608.

5. Xu J. Preparation, culture, and immortalization of mouse embryonic fibroblasts. *Curr Protoc Mol Biol*. 2005;Chapter 28:Unit 28 1.

6. Ackers-Johnson M, Li PY, Holmes AP, O'Brien SM, Pavlovic D and Foo RS. A Simplified, Langendorff-Free Method for Concomitant Isolation of Viable Cardiac Myocytes and Nonmyocytes From the Adult Mouse Heart. *Circ Res*. 2016;119:909-20.

7. Zhang H, Pan B, Wu P, Parajuli N, Rekhter MD, Goldberg AL and Wang X. PDE1 inhibition facilitates proteasomal degradation of misfolded proteins and protects against cardiac proteinopathy. *Sci Adv*. 2019;5:eaaw5870.

8. Chen Q, Liu JB, Horak KM, Zheng H, Kumarapeli AR, Li J, Li F, Gerdes AM, Wawrousek EF and Wang X. Intrasarcoplasmic amyloidosis impairs proteolytic function of proteasomes in cardiomyocytes by compromising substrate uptake. *Circ Res*. 2005;97:1018-26.

9. Pan B, Li J, Parajuli N, Tian Z, Wu P, Lewno MT, Zou J, Wang W, Bedford L, Mayer RJ, Fang J, Liu J, Cui T, Su H and Wang X. The Calcineurin-TFEB-p62 Pathway Mediates the Activation of Cardiac Macroautophagy by Proteasomal Malfunction. *Circ Res*. 2020;127:502-518.

10. Guo L, Prall W and Yang X. Assays for the Degradation of Misfolded Proteins in Cells. *J Vis Exp*. 2016.

11. Ranek MJ, Terpstra EJ, Li J, Kass DA and Wang X. Protein kinase g positively regulates proteasome-mediated degradation of misfolded proteins. *Circulation*. 2013;128:365-76.
